# Rapid nuclear exclusion of Hcm1 in aging *Saccharomyces cerevisiae* leads to vacuolar alkalization and replicative senescence

**DOI:** 10.1101/268664

**Authors:** Ata Ghavidel, Kunal Baxi, Martin Prusinkiewicz, Cynthia Swan, Zach R. Belak, Christopher H. Eskiw, Carlos E. Carvalho, Troy A. Harkness

## Abstract

Yeast cells, like other higher eukaryotic cells, undergo a finite number of cell divisions before exiting the cell cycle due to the effects of aging. Here, we show that yeast aging begins with the nuclear exclusion of Hcm1 in young cells, resulting in loss of acidic vacuoles. Autophagy is required for healthy aging in yeast, with proteins targeted for turnover by autophagy directed to the vacuole. Consistent with this, vacuolar acidity is necessary for vacuolar function and yeast longevity. Using yeast genetics and immunofluorescence microscopy, we confirm that vacuolar acidity plays a critical role in cell health and lifespan, and is potentially maintained by a series of Forkhead Box (Fox) transcription factors. An interconnected transcriptional network involving the Fox proteins (Fkh1, Fkh2 and Hcm1) are required for transcription of v-ATPase subunits and vacuolar acidity. As cells age, Hcm1 is rapidly excluded from the nucleus in young cells, blocking the expression of Hcm1 targets (Fkh1 and Fkh2), leading to loss of v-ATPase gene expression, reduced vacuolar acidification, increased α-syn-GFP vacuolar accumulation, and finally, diminished replicative lifespan (RLS). Loss of vacuolar acidity occurs about the same time as Hcm1 nuclear exclusion and is conserved; we have recently demonstrated that lysosomal alkalization similarly contributes to aging in *C. elegans* following a transition from progeny producing to post-reproductive life. Our data points to a molecular mechanism regulating vacuolar acidity that signals the end of RLS when acidification is lost.

## Introduction

The Hayflick Limit describes how proliferating eukaryotic cells undergo a finite number of cell divisions before succumbing to aging phenotypes (Hayflick and Moorhead 1961; Hayflick 1965; Burnet 1974), such as alterations to cell morphology, genome architecture, and organelle functions (Cuervo and Dice 2000; Hwang *et al.* 2009; Feser *et al.* 2010). The effects of aging do not appear to simply be a default outcome of general cell deterioration, but may instead be actively regulated via genetically controlled longevity pathways (Kaeberlein et al. 2005; DiLoreto and Murphy 2015). The simple brewing yeast, *Saccharomyces cerevisiae,* has provided a wealth of information regarding the genetic nature of lifespan determination (Arlia-Ciomo *et al.* 2014; Swinnen *et al.* 2014; Wierman and Smith 2014). The value of using yeast to study longevity becomes clear when the conservation of the longevity pathways, from yeast to humans, is considered (Mirzaei *et al.* 2014; DiLoreto and Murphy 2015; Ruetenik and Barrientos 2015). Current thought supports the idea that nutrient and stress response pathways play antagonistic roles in maximizing cell health. According to the Hormesis theory, small stresses promote longevity by activating signalling networks that increase cell repair and slow cell growth (Shore and Ruvkun 2013). On the other hand, nutrient response pathways, primarily consisting of the mTOR/AKT/Insulin signalling networks (Sch9/Tor1/Snf1 in yeast), shunt resources into utilizing food when it is available. These pathways run opposed to hormetic pathways, as mutations to nutrient response pathways lead to increased stress resistance and prolonged longevity (Swinnen *et al.* 2014; Hu et al. 2014).

Stress in yeast is managed by several conserved families of proteins that form highly integrated transcriptional networks. The Forkhead Box (Fox) proteins in higher eukaryotes, such as the FOXO class of proteins, are tightly correlated with stress response and tumor suppression (Chiacchiera and Simone 2010; Martins *et al.* 2016). In *C. elegans*, it is firmly established that DAF-16, the sole FOXO orthologue in worms, is critical for normal lifespan, and lifespan extension when the insulin-signalling pathway is shut down (Murphy 2013; Antebi 2013). It is now apparent that the yeast redundant Fox proteins Fkh1 and Fkh2 (both *FKH1* and *FKH2* must be deleted to observe a phenotype) are also critical for stress response and lifespan (Zhu *et al*. 2000; Shapira *et al.* 2004; Postnikoff *et al.* 2012; Linke *et al.* 2013; Jiao *et al.* 2015; Malo *et al.* 2016). A third yeast Fox family member, Hcm1, controls Fkh1 and Fkh2 transcription during G2, which regulates cell cycle progression (Pramila *et al*. 2006). Alternatively, under stress conditions, these three Fox proteins work in a positive feedback loop with the Snf1 kinase, a metabolic stress response factor orthologous to the mammalian AMP-activated protein kinase (AMPK; Hedbacker and Carlson 2008; Ghillebert *et al.* 2011; Rodriguez-Colman *et al.* 2013; Jiao *et al.* 2015). When activated by stress, Snf1 phosphorylates Hcm1, driving it into the nucleus where it transcribes its target genes, including *FKH1* and *FKH2* (Rodriguez-Colman *et al.* 2013). Fkh1 and Fkh2 then reinforce Snf1 activity by transcribing *SNF1* (Jiao *et al.* 2016). Thus, a series of highly conserved stress responsive signalling pathways are intertwined in yeast to tightly regulate changes in gene expression and impact longevity.

As cells age, proteotoxic stress mechanisms can no longer cope with accumulating cellular damage, leading to increased protein aggregation (Tenreiro *et al.* 2013; Kim *et al.* 2016; Kikis 2016). While protein aggregation in aging mammalian cells is linked with neurodegenerative disease, it may also provide an adaptive mechanism to protect proteins from stress and the effects of aging (Miller *et al.* 2015; Saarikangas and Barral 2016). However, mechanisms facilitating proteostasis as cells age remain unclear. In yeast, it has been shown that protein aggregates are asymmetrically inherited during cell division, such that mother cells retain the bulk of the damaged proteins via a retention mechanism consisting of heat shock proteins and cytoskeletal elements (Erjavec *et al.* 2007). Asymmetric inheritance in yeast ensures daughter cells are born with the best chance at a full lifespan, and also extends to vacuoles, the end-point of proteolytic breakdown of damaged and misfolded proteins. Vacuolar acidity facilitates the proper activity of vacuolar enzymes, and is renewed in daughter cells, but not in mother cells (Henderson *et al.* 2014), thus ensuring daughters are born with fully functional acidic vacuolar compartments. It has been shown in yeast that vacuolar acidity is linked with both extended replicative lifespan (Hughes and Gottschling 2012; Henderson *et al.* 2014) and chronological lifespan (Ruckenstuhl et al. 2014). It is currently believed that loss of vacuolar acidity in aging cells leads to cellular impairment and senescence, and may be due to mitochondrial dysfunction (Ohya *et al.* 1991; Merz and Westermann 2009; Hughes and Gottschling 2012). Nonetheless, it remains unresolved whether impaired proteolytic function in alkalizing vacuoles is a driving force in aging. Recent literature, however, links the integrative stress response in yeast with enhanced replicative lifespan and autophagy (Postnikoff *et al.* 2017; Tyler and Johnson 2018). To address the question of whether proteolytic dysfunction in old, alkalized vacuoles plays a role in aging, we monitored the proteolytic degradation of a human protein in aging yeast cells that forms inclusions in patients with a variety of neurodegenerative diseases (α-synuclein; Yang and Yu 2016) and observed that aggregates accumulated within vacuoles as cells age. We show that aggregates accumulate as vacuoles alkalize, and that enhanced α-synuclein aggregation decreases RLS. Our observations support the idea that maintenance of vacuolar acidity is a vital contributor to prolonged lifespan and that the Fox proteins, Fkh1, Fkh2 and Hcm1, play an important role in regulating vacuolar alkalization. Our recent work demonstrates that this is an evolutionarily conserved molecular mechanism from yeast to worms (Baxi *et al.* 2017).

## Materials and Methods

### Yeast strains and methods

Unless indicated otherwise, experiments were carried out using the BY4741 background (*MAT***a** *his3Δ1 leu2Δ0 met15Δ0 ura3Δ0*). Respective null mutants were constructed by single step PCR-based gene deletion. Strains were grown in rich (yeast/peptone; YP) with the appropriate sugar supplemented (i.e. glucose or galactose) or defined (synthetic complete; SC) media supplemented with 30 μg/ml of all amino acids. Yeast transformations, *E.coli* DNA extractions, and flow cytometry were done as previously described (Malo *et al.* 2016).

### Replicative Lifespan analysis of yeast

RLS assays were done essentially as described (Postnikoff and Harkness 2014). Briefly, cells from logarithmically growing liquid cultures were struck out onto YPD plates. After an overnight incubation at 30°C, a minimum starting population of 40 newly budded cells were removed to start the experiment using a Zeiss Micro-manipulator, where the new buds would serve as the mother cells. Buds were successively dissected away until all mother cells had ceased dividing. The plates were maintained at 4°C overnight.

### Old yeast cell enrichment, RNA preparation and analysis

Yeast cultures enriched for cells in old replicative stages were prepared following the protocol for the Mother Enrichment Program (MEP) described elsewhere (Lindstrom and Gottschling 2009. Yeast RNA was prepared by TRIZOL (Life Technologies) extraction. Following extensive DNase I treatment, first strand cDNA was synthesized using an oligo(dT) primer and reverse transcriptase (Fermentas) at 42°C for 1 h. Query RNAs were amplified at incremental PCR cycles using RNA specific primers and resolved on agarose gels containing ethidium bromide. Quantitative PCR was carried out in tandem using a real-time cycler and iQ SYBR Green qPCR SuperMix (Bio Rad). Assays were done in triplicates. When noted, two independent RNA preparations were analyzed. A single melt peak confirmed the identity of each PCR product. Each assay included a no-template control for every primer set. Primer sequences are available on request. MEP was used for RNA analyses and to generate data for the plot shown in Fig. 1B.

**Figure 1.**
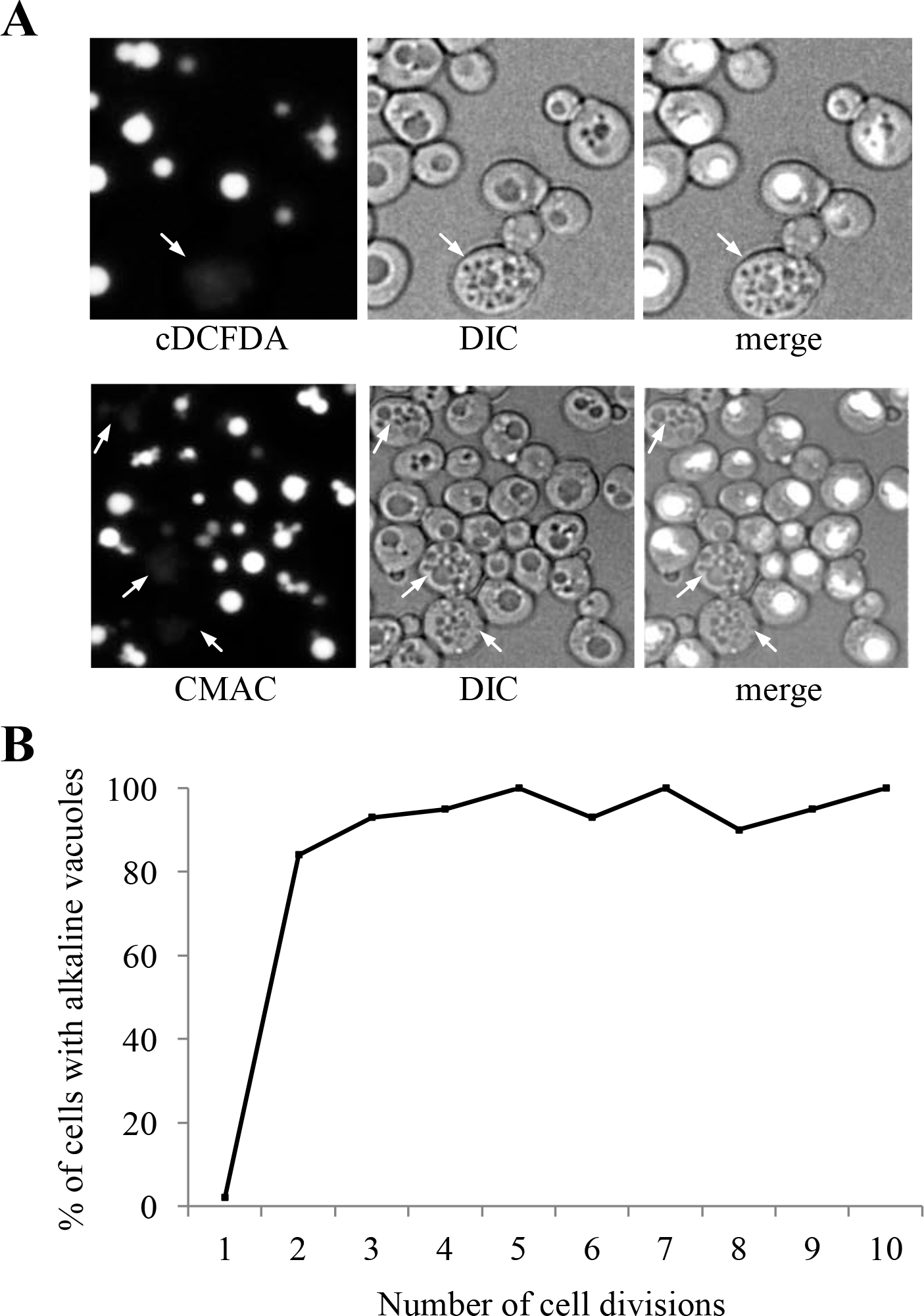
Vacuolar morphology and acidity change as yeast cells age. **(A)** WT cultures grown overnight were stained with the pH sensitive vacuolar fluorescent probes CMAC-Arg or carboxy-DCFDA (cDCFDA). Both stains fluoresce when subjected to an acidic environment. Cells were grown and stained at physiological pH. Arrows denote representative older cells with fragmented vacuoles that lack staining, indicative of alkalized vacuoles. See Supplemental Figs. 1A and B showing that yeast cells with greater than 4 buds scars typically display fragmented vacuoles. (**B**) Populations were enriched for older cells using the Mother Enrichment Program (MEP; Lindstrom and Gottschling 2009). Cells were binned according to their indicated number of bud scars and scored as having alkaline vacuoles if a 2-fold or greater reduction in fluorescence intensity was observed compared to young cells with one bud scar. Note that the plot reaches a steady state in cells with greater than 3 divisions. The bins contain anywhere between 20 (for cells with high number of bud scars) and 40 (for cells with few number of scars) cells. See Supplemental Fig. 1C for cDCFDA fluorescence emission spectra of young and old yeast.

### Preparation and analysis of yeast protein lysate

Total yeast protein lysates were prepared and analyzed as described (Ghavidel *et al.* 2007). Mouse α-GFP (Covance) and rabbit α-TAP (Pierce) antibodies were used at 1/5000.

### Staining yeast vacuoles

Staining with vacuolar probes CMAC-Arg (7-amino-4-chloromethylcoumarin, L-arginine amide) and carboxy-DCFDA (5-carboxy-2,7, dichlorofluorescein diacetate) were performed according to manufacturer’s instructions (Molecular Probes). Briefly, cells were treated with 100 uM CMAC-Arg or 10 uM cDCFDA in the same growth media (at physiological pH) for approximately 1 hour prior to imaging. cDCFDA fluorescence results from diffusional uptake and subsequent hydrolysis by vacuolar esterase. cDCFDA is non-fluorescent until the acetate groups are cleaved to yield the fluorescent fluorophore, 5-(and-6)-carboxy-2′,7′ –dichlorofluorescein (Preston et al. 1989). The hydrolysis product is a relatively impermeable anion and is trapped in cellular compartments where it forms. With a pK_a_ of 4.8, this dye is used as an acidic pH sensor. We maintained cells at physiological pH when using cDCFDA. Cells maintained at pH 4 resulted in staining of vacuoles in older cells (Supplemental Fig. 1E; Vida and Emr 1995); cells grown in acidic media likely have a lower vacuolear pH than those grown at the physiological pH.

### Native ChIP

Cells from 50 ml early log cultures were lysed in buffer T (20 mM Tris-HCl (pH 8), 80 mM KCl, 0.1% TX100, 1 mM DTT, protease inhibitors) by bead beating for 20 sec. The slurry was sonicated on ice and spun at 5,000 rpm for 10 minutes. TAP complexes were immunoprecipitated from the supernatant using an α-TAP rabbit polyclonal antibody (Abcam), washed extensively in lysis buffer, and eluted in 0.1 M Glycine (pH 1.3) at RT. Eluates were neutralized with 1/5 V:V 1.5 M Tris-HCl (pH 8.8) and incubated with 2 μg/μl DNase-free RNase A (Fermentas) at 37°C for 30 minutes. Recovered protein-DNA complexes were deproteinized by incubating with 2 μg/μl Proteinase K (Thermo Fisher) at 42°C for 30 minutes, extracted with phenol chloroform, supplemented with 1 μg glycogen as carrier, and ethanol precipitated overnight at 20°C. Pellets were washed in 70% ethanol and resuspended in nuclease free water. Genomic DNA was amplified using primers to 5’ upstream regions of *VMA* genes. *VMA6* (YLR447C) encoding a V_0_ integral membrane subunit of V-ATPase harbors a putative Fkh binding site. *VMA7* (YGR020C) encodes a V_1_ peripheral membrane subunit of V-ATPase and its promoter region does not display an obvious consensus binding motif for Fkh. Wild type (WT) cells, which do not contain a TAP fusion, were IPed in parallel as experimental controls. As an additional control, a α-GST rabbit polyclonal antibody (Abcam) was used for IP when indicated.

### α-Syn and vacuolar imaging

Yeast cells were spotted on a glass slide and imaged with an Olympus BX-51 microscope equipped with Infinity software v.5.0.3 (Lumenera, Ottawa, Canada) for image acquisition. Alternatively, mounted cells were imaged using a laser scanning confocal microscope (Zeiss LSM 510). MetaMorph v6.1 software (Universal Imaging Corporation; Downington, PA) was used to reconstruct 3D images from the calibrated overlays of the z-stacks. Emission spectra were obtained by scanning sample fluorescence intensity over an emission wavelength of 400-700 nm while applying a constant excitation wavelength of 495 nm (Preston et al. 1989). To image old cells, calcofluor-white, which stains chitin in bud scars was used (Bartholomew and Mittwer 1953; Cabib and Bowers 1971; Smeal et al. 1996).

### Hcm1-GFP imaging

Hoechst and calcofluor-white are imaged in the same channel and therefore cannot be used together. Thus, to image nuclei and bud scars in the same cell, we used Wheat Germ Agglutinin–Alexafluor 555 conjugate (WGA-AF555; Invitrogen (Cat. #W32464)). WGA-AF555 was dissolved in PBS at a concentration of 1 mg/mL and stored at −80 C until use. Hoescht 33342 fluorescent DNA stain was obtained as a 10 mg/mL solution in water from Life Technologies Inc. (Cat. #H3570), stored at 4 C, and diluted to 1 mg/mL in water before use. Yeast cells were grown to mid-log phase in Synthetic Complete Medium (YPD tends to cause a high level of background fluorescence) containing 2% w/v glucose. Cells were fixed by addition of 4% final concentration of formaldehyde and incubation at room temperature for 1 h. Cells were collected by centrifugation (1500 X G, 5 min), and washed twice in water. Aliquots of cells were spun down and resuspended in 200 μL of 0.2 M Sodium dihydrogen orthophosphate and treated for 20 min on ice with 2 μL of a 100 mg/mL solution of sodium borohydride in 14 M NaOH. Cells were diluted with 1 mL of 100 mM Tris-HCl pH 7.5, collected by centrifugation and washed twice more with 1 mL 100 mM Tris-HCl pH 7.5. Cells were then suspended in 1 mL 100 mM Tris-HCl pH 7.5 and treated with 1 μL of 1 mg/mL Hoescht 33342 (1 μg/mL final) and 5 μL of 1 mg/mL WGA-AF555 (5 μg/mL final concentration), followed by incubation with gentle agitation at room temperature for 30 min in the dark. Cells were again collected by centrifugation, pellets resuspended in 100 μL of solid aqueous mounting medium pre-heated to 38°C and applied to warmed slides followed by addition of coverslips. Medium was permitted to harden for 1 hr in the dark before imaging of cells. All cells were imaged using identical exposure times in the GFP, AF555, and H33342 channels. Cells were initially imaged in the AF555 channel to count bud scars and then subsequently imaged in the GFP and H33342 channels. ImageJ was used to create masks based on the GFP signal channel and pixel intensity within the mask was summed to give total cellular GFP signal. Similarly, masks were created using the H33342 signal and applied to the GFP images and pixel intensity with the mask area was summed to give total nuclear GFP signal. Percent nuclear GFP was calculated for each selected cell individually and average nuclear signal fraction averaged. Statistics used a student’s two-tailed t-test to compare ascertain the significance of the observed reductions in nuclear GFP signal.

### Image quantification and data analyses

Relative fluorescence intensity values were calculated using the multi-platform open source software ImageJ. Fluorescence values were exported to Microsoft Excel and subsequently plotted in box-and-whisker format using R software. Statistical analysis was performed using GraphPad Software’s online statistical calculator available at https://www.graphpad.com/quickcalcs/ttest2/. See Supplemental Tables 2-5 for a complete analyses fo each cell imaged and for a summary of the calculated p-values.

## Results

### Vacuolar fragmentation and loss of vacuolar acidity in aging yeast cells

Proteolytic degradation in vacuoles is the primary method for removal of protein aggregates not amenable to refolding by molecular chaperones or degradation by the proteasome (Marques *et al*. 2006). Vacuoles are membrane-bound organelles that contain an assortment of acid-dependent hydrolases. The acidic lumen of vacuoles, critical for enabling proteolytic degradation, is maintained by the activity of a vacuolar proton-coupled ATPase (v-ATPase), conserved across evolutionary boundaries (Nelson 1992; Preston et al. 1992). As described above, loss of vacuolar acidity has been observed in aging yeast cells using the vacuolar pH sensitive monitors quinacrine (decreasing fluorescence as vacuoles alkalize) and Pho8-super ecliptic pHluorin (Pho8-SEP; increasing fluorescence as acidity is lost; Hughes and Gottschling 2012; Henderson et al. 2014). We show that aging cells do indeed suffer vacuolar fragmentation as early as after 4 doublings, which is visible using DIC imaging (Supplemental Fig. 1A) or fluorescent imaging when using the vacuolar membrane stain FM4-64 (Supplemental Fig. 1B; Cole *et al.* 1998; Wiltshire and Collings 2009). We used three additional methods to measure vacuolar acidity: the pH sensitive vacuolar dyes cDCFDA (Preston *et al.* 1989; Baxi *et al.* 2017) and CMAC-Arg (Roberts *et al.* 1991), and the Ste3-GFP and Ste3-pHluorin fusion constructs. Both CMAC and cDCFDA stain acidic compartments in cells with fragmented vacuoles (Fig. 1A; cDCFDA staining quantified in Supplemental Fig. 1C). cDCFDA also stains acidic lysosomes in intestines of live *C. elegans* (Baxi *et al.* 2017). In Matα yeast cells, Ste3-GFP is targeted to the vacuole for degradation where the GFP moiety remains stably detectable for extended periods of time. The GFP variant, pHluorin, also remains detectable following Ste3-pHluorin degradation in the vacuole, and exhibits increased fluorescence as the pH of the intracellular environment rises (Supplemental Fig. 1D; Miesenbock *et al.* 1998). As a further control for cDCFDA acidic-dependent fluorescence, we grew cells in YPD media maintained at pH 4, which is sufficient to acidify vacuoles through passive diffusion (Nelson and Nelson 1990). After 6 hours of growth in acidic media, cDCFDA fluorescence was comparable in vacuoles of young and older mother cells (Supplemental Fig. 1E). Using calcofluor-white, which stains chitin, a substance that is concentrated in bud scars (Bartholomew and Mittwer 1953; Cabib and Bowers 1971; Smeal et al. 1996), cell age can be estimated. We demonstrated using these approaches that vacuole acidity does indeed decrease rapidity as yeast cells age (Fig. 1B).

### Reduced v-ATPase expression and function is associated with reduced lifespan

We measured RNA levels of genes involved in vacuolar maintenance and function in a population enriched for replicatively old yeast cells (>7 divisions), and observed that expression of multiple vacuolar proton pump subunits was reduced compared to WT (Fig. 2A; quantified in Supplemental Fig. 2A). In this population of enriched cells, we could isolate enough RNA to perform RT-PCR, but unfortunately could not isolate enough protein for western analyses. As controls, we measured the expression of *YPT7* and *SIP18*, which encode a protein required for homotypic fusion of vacuolar membranes, and an osmotic stress protein elevated in older cells, respectively, and found that *YTP7* expression was reduced and *SIP18* expression was elevated in older cells (Fig. 2A; quantified in Supplemental Fig. 2A). This supported findings by us (Fig. 1; Supplemental Fig. 1) and others (Tang et al. 2008; Gebre et al. 2012) that vacuolar fragmentation occurs in old cells.

**Figure 2.**
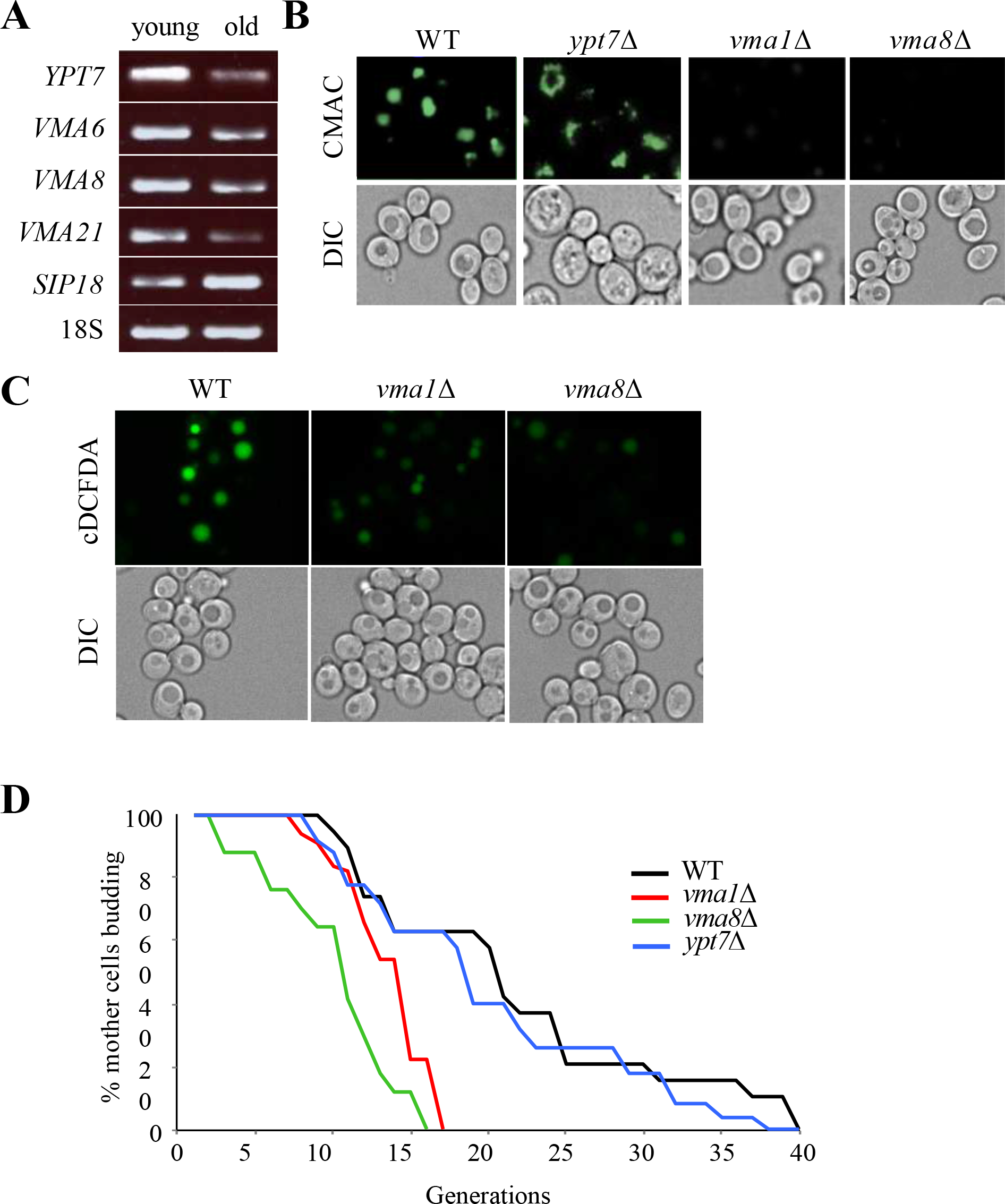
The yeast v-ATPase is required for acidity of the vacuolar lumen and full replicative lifespan. **(A)** RT-PCR RNA expression analyses of vacuolar proton pump subunits (*VMA*) and a vacuolar fusion factor (*YPT7*) in replicatively old cells enriched using MEP. Agarose gel electrophoresis of ethidium bromide stained RT-PCR products is shown. Sip18, encoding an osmotic stress protein previously reported to be upregulated in old cells (Lesur and Campbell 2004), served as a control. See Supplemental Fig. 2A for quantification of old:young mRNA ratios. **(B)** CMAC-Arg fluorescence imaging of isogenic dividing WT and single deletion yeast mutants. Cells were stained, processed, and imaged in parallel. See Supplemental Fig. 2B for CMAC-Arg fluorescence emission spectra of cells shown here. **(C)** cDCFDA fluorescence imaging of isogenic dividing WT and single deletion yeast mutants. Cells were stained, processed, and imaged in parallel. **(D)** RLS of WT yeast and deletion mutants on YPD media. Statistical analysis is available in Table S1.

To determine whether the loss of vacuolar acidity is an end result of cellular aging, or if it actively drives the cell into replicative senescence, we examined vacuolar acidity and replicative longevity in cells harbouring deletions of *YPT7*, which exhibit fragmented vacuoles, or deletions of genes encoding the v-ATPase subunits *VMA1* and *VMA8*. Cells lacking *YPT7* maintained both WT vacuolar acidity and normal RLS (Figs. 2B and 2D; Table S1), despite their extensively fragmented vacuoles. In contrast, *vma1*Δ and *vma8*Δ cells contained alkalized vacuoles, measured using CMAC (Fig. 2B) and cDCFDA (Fig. 2C), and a correspondingly reduced RLS (Figs. 2D; Table S1). The loss of RLS in v-ATPase mutants highlights a requirement for vacuolar acidity in extending longevity in yeast.

### Accumulation of α-synuclein-GFP in vacuoles of aging yeast cells is linked to shortened lifespan

In older replicating WT cells, irreversibly misfolded and damaged proteins progressively sequester into insoluble high molecular weight cytoplasmic aggregates (Erjavec et al. 2007; David et al. 2010). Once transported to vacuoles by autophagy, protein aggregates are cleared by the proteolytic activity of hydrolases (Cuervo et al. 2005). Vacuolar hydrolases function optimally over a narrow range of low pH (Bachhawat et al. 1993), and as a consequence, are sensitive to subtle perturbations in pH. We reasoned that the loss of vacuolar acidity in older yeast cells could lead to hydrolase dysfunction, and failure to clear protein aggregates in these organelles. To test this idea, we monitored steady state levels of GFP alone, or GFP-tagged human α-synuclein (α-syn-GFP) expressed from a *GAL1* promoter in dividing yeast cultures (Outeiro and Lindquist 2003). Formation of α-syn aggregates coincides with the onset of human neurodegenerative disorders broadly associated with aging (Muchowski 2002). GFP is expressed in the cytoplasm of young and old cells, yet is distinctly excluded from vacuoles (Fig. 3A, top panels). Newly expressed α-syn-GFP in vegetatively growing cells was initially localized to the plasma membrane of daughter cells, due to its affinity for phospholipids (Fig. 3A, middle panels; Dixon *et al.* 2005; Popova *et al.* 2015). In WT mother cells, on the other hand, α-syn-GFP aggregates began to accumulate as cytosolic inclusions (Fig. 3A, middle panels). Increased expression of α-syn-GFP from the *GAL* promoter ultimately resulted in larger cytoplasmic inclusions than when expressed at lower levels (Supplemental Fig. 3A; Petroi *et al.* 2012). α-syn-GFP is normally degraded by the ubiquitin-proteasome system, but larger aggregates are instead cleared via autophagy (Ebrahimi-Fakhari *et al.* 2011; Xilouri *et al.* 2013). Consistent with this, very few dividing cells expressed an intravacuolar GFP signal (2.8%, n=38; Fig. 3A, middle panels). However, in *vma8*Δ cells with alkalized vacuoles, 43% (n=42) of the cells expressed α-syn-GFP vacuolar aggregates (Fig. 3A, lower panels). To determine whether increased vacuolar aggregates were in fact due to decreased vacuolar proteolysis, we used antibodies against GFP to visualize α-syn-GFP cleavage products in protein lysates of WT and *vma8*Δ cells. Compared to WT cells, *vma8*Δ cells harbored a larger pool of uncleaved α-syn-GFP (Fig. 3B). We also observed the localization of α-syn-GFP predominantly in vacuoles of young dividing cells lacking the protease Pep4, similar to that documented in *vma8*Δ cells (92%, n=25; Fig. 3C), consistent with the notion that vacuolar proteolysis loses its effectiveness as cells age. The cytoplasmic α-syn-GFP inclusions commonly observed in young dividing WT cells were no longer visible in older cells, but rather, were viewed as vacuolar inclusions (Fig. 4A).

**Figure 3.**
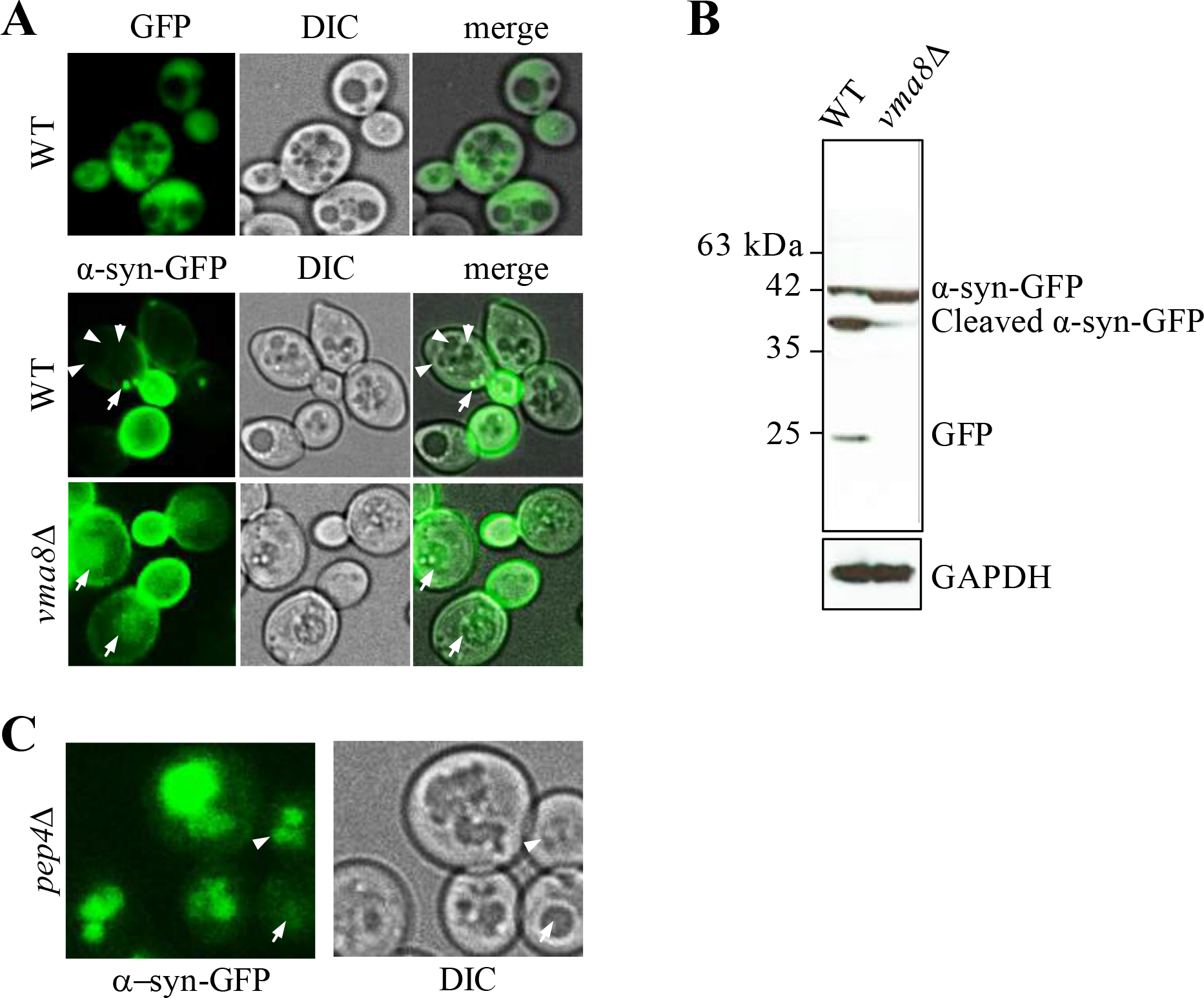
Cells with impaired vacuolar function accumulate uncleaved α-syn-GFP aggregates within their vacuoles. **(A)** Live cell confocal imaging of genomically integrated α-syn-GFP expressed from a *GAL1*-inducible promoter in WT and *vma8Δ* mutants grown in YP + 0.1% galactose. Expression of a GFP reporter using the *GAL1* promoter results in cytoplasm fluorescence (top panel). Arrows denote cytoplasmic GFP foci and arrowheads point to vacuolar lumen in WT cells. See Supplemental Fig. 3A for images of α-syn-GFP aggregation as galactose concentrations increase. **(B)** GFP westerns in whole cell lysates prepared from cells in **(A)**. GAPDH served as a loading control. **(C)** Live cell imaging of dividing *pep4*Δ cells expressing genomically integrated α-syn-GFP induced from a *GAL1* promoter. Cells were grown in 0.1% galactose. The arrow depicts a very young cell with subtle vacuolar staining; the arrowhead shows a young cell experiencing vacuolar fragmentation with marked vacuolar staining.

Representative images of the cells binned into 3 age groups are shown. Accumulation of α-syn-GFP in aging cells was indeed occurring within alkalizing vacuoles, since α-syn-GFP and reduced CMAC staining colocalize only in older cells (Fig. 4B). Ratios of vacuolar:cytosolic fluorescent staining from the cells binned in Fig. 4A were determined (Fig. 4C), and demonstrate that a shift towards vacuolar localization occurs early, consistent with vacuolar alkalization (Fig. 1B). The ongoing efficiency in aging cells regarding relocalization of α-syn-GFP aggregates from the cytosol to the vacuole suggests that the formation, transport, and docking of autophagy vesicles to vacuoles likely remains intact in aging cells, an observation supported by the increased induction of autophagy (ATG) genes in these cells (Supplemental Fig. 3B, quantified in lower panel).

**Figure 4.**
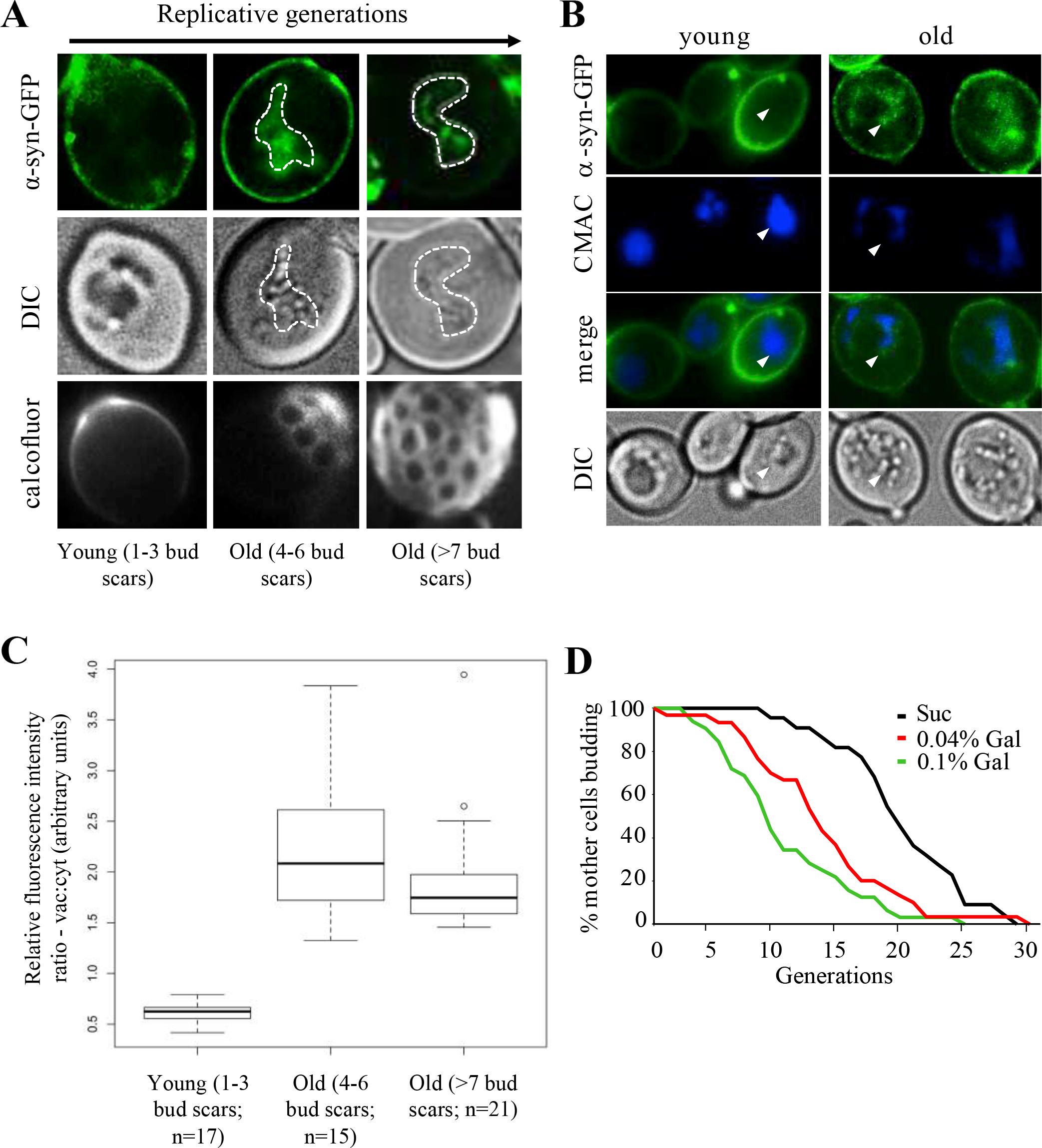
Accumulation of α-syn-GFP protein aggregates in replicatively old yeast. **(A)** Live single cell confocal imaging of α-syn-GFP in dividing WT yeast cultures. Calcofluor stained bud scars are shown in bottom panels. Vacuole boundaries are marked by white dashed lines. **(B)** Intravacuolar compartmentalization of protein aggregates in replicatively old yeast. Cells expressing α-syn-GFP were grown in 0.1% galactose and counter stained with CMAC-Arg for 10 minutes prior to imaging. Arrows denote vacuolar boundaries. **(C)** Box & whisker plot of vacuolar:cytoplasmic GFP fluorescence ratios in cells undergoing early or late divisions. Digitized fluorescence images were scored by an observer blind to cell age. Mean, and upper and lower quartiles are included. The cells were grouped into 3 age groups, as shown, with the specific (n)s for each group indicated. **(D)** RLS of WT yeast expressing *GAL1*-inducible α-syn-GFP grown on YP media supplemented with sucrose (α-syn-GFP “off’) or galactose (α-syn-GFP “on”). Mean lifespan for a population of 40 mother cells is: Suc, 18.2; 0.04% Gal, 12.5; 0.1% Gal, 8.1.

A prediction based on these observations is that the loss of vacuolar acidity, resulting in the accumulation of vacuolar protein aggregates in older dividing cells, contributes to replicative senescence. To test this idea, we monitored RLS in yeast that expressed *GAL* inducible α-syn-GFP. Higher expression of α-syn-GFP resulted in a dose-dependent reduction in median RLS (Fig. 4D). Although this observation is consistent with the literature, as formation of protein aggregates in post mitotic cells is associated with aging (David *et al.* 2010; Erjavec *et al.* 2007), conclusions must be made cautiously, as α-syn-GFP expression in yeast has been associated with numerous phenotypes, including induction of oxidative stress, and mitochondrial dysfunction (Popova *et al.* 2015), any of which could impact yeast RLS.

### Fhk1 or Fkh2 are required for regulated vacuolar acidification

Loss of vacuolar acidity is clearly a hallmark of aging yeast cells and may ultimately influence RLS. Understanding how to prevent aging-related vacuolar alkalization may therefore enhance RLS. The simultaneous transcriptional downregulation of *VMA* genes in aging cells (Fig. 2A) suggested that a common regulatory mechanism may control expression of this gene set, with potentially a discrete set of transcription factors involved. Analyses of *VMA* promoter regions revealed putative binding motifs for members of the Forkhead Box (Fox) family of transcription factors (Supplemental Fig. 4). The homologous yeast Fox proteins, Fkh1 and Fkh2, drive the cyclical expression of genes required for G2/M progression in yeast, and are for the most part functionally redundant (Zhu *et al.* 2000; Hollenhorst *et al.* 2000). Using a native chromatin immunoprecipitation (ChIP) assay, we demonstrated that Fkh1 and Fkh2 bind to the *VMA6* and *8* promoters with comparable affinity (Fig. 5A). As a control, we show that the *VMA7* promoter, which does not contain a Fox consensus site, did not bind Fkh1 or Fkh2 (Fig. 5A). Our results are consistent with a previous report of Fkh1/Fkh2 binding sites within the *VMA3*, *5*, *9* and *13* promoters (Ostrow *et al.* 2014). Consistent with their overlapping functional roles, deleting either *FKH1* or *FKH2* conferred no obvious defect in vacuolar acidity (Fig. 5B). Deleting both *FKH1* and *FKH2*, however, led to loss of vacuolar acidity (Fig. 5B) and reduced expression of *VMA* genes in young dividing cells (Fig. 5C). The downregulation of *VMA* genes in *fkh1Δ fkh2Δ* cells was subtle in magnitude (between 30 to 50% reduction), yet broad in scope (Fig. 5C), and similar to that observed in older cells (Fig. 2A). The decrease in both *FKH1* (30%) and *FKH2* (50%) mRNA expression in older cells may explain the reduced expression of *VMA* genes (Fig. 5D). Thus, downregulation of Fkh-mediated transcription in aging yeast cells provides a testable mechanism to explain the loss of vacuolar acidity in old yeast.

**Figure 5.**
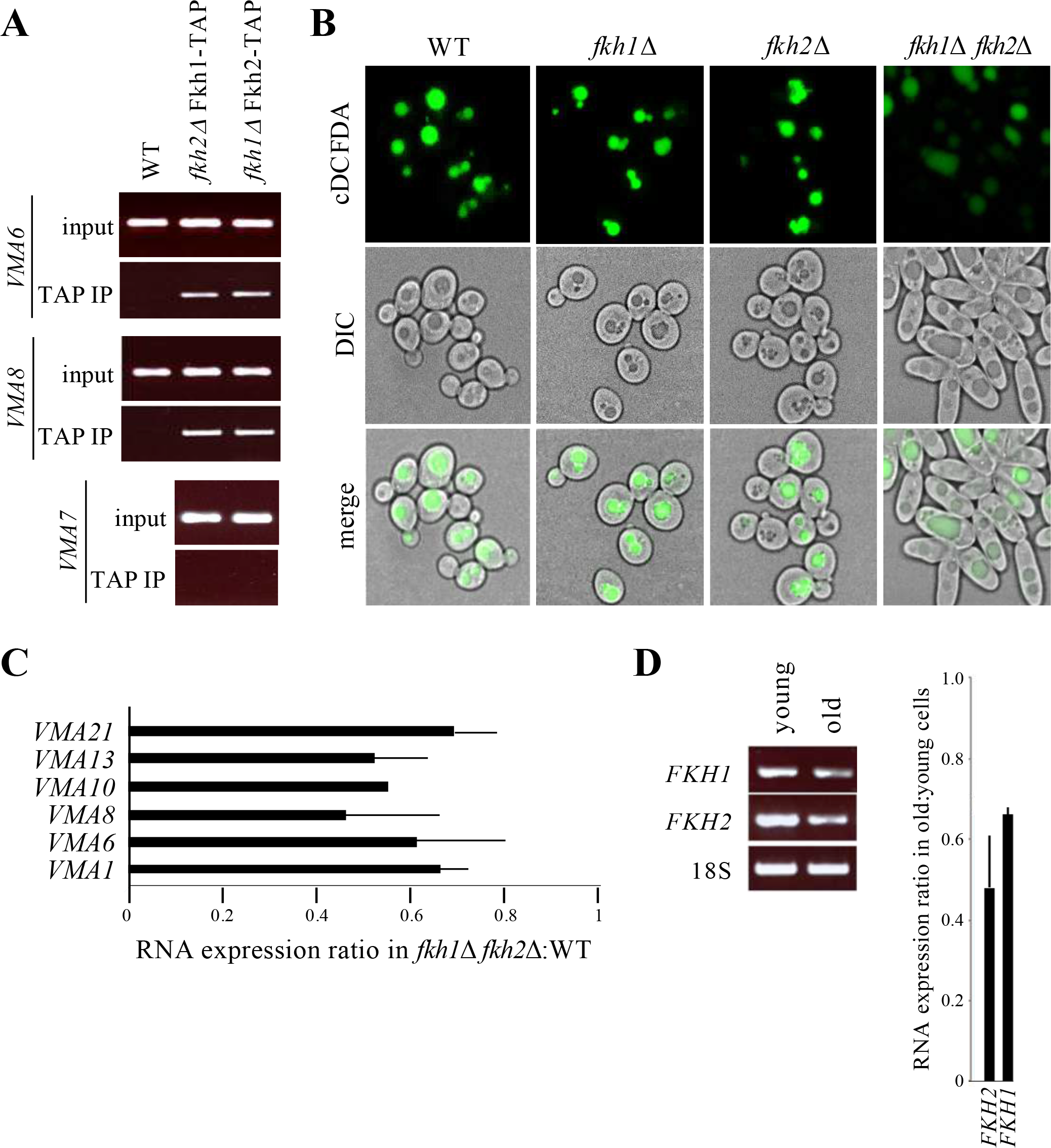
Vacuolar acidity is controlled by Fkh1 or Fkh2. **(A)** Native chromatin immunoprecipitation (ChIP) of select *VMA* promoters in cells expressing the TAP epitope fused to the C-terminus of *FKH1* or *FKH2* (Postnikoff *et al.* 2012). Genomic DNA in TAP immunoprecipitates was amplified using primers to 5’ upstream regions. See Supplemental Fig. 4 for *FKH* consensus binding sites within *VMA* promoters. **(B)** Mutants harboring deletions of either or both *FKH* genes were grown overnight in YPD, then stained with cDCFDA and imaged in parallel. Deleting both *FKH* genes induced nutrient-independent pseudohyphal growth (Zhu *et al.* 2000; Hollenhorst *et al.* 2000). **(C)** qPCR analyses of *VMA* mRNA expression ratios in *fkh1Δ fkh2Δ* mutants relative to WT cells. Mean ± SD shown. **(D)** RT-PCR analyses of *FKH* mRNA in young and old cells enriched by MEP. Expression ratio of *FKH* genes in old cells relative to young cells. Mean ± SD shown.

### Vacuolar alkalization coincides with nuclear exclusion of Hcml in old cells

*FKH* gene expression is activated in a cell cycle dependent manner in G2 by a third Fox protein, Hcm1 (Pramila *et al.* 2006). Yeast *HCM1* encodes a transcription factor involved in chromosome segregation, spindle assembly and budding, and is transcribed at the G1/S boundary (Horak *et al*. 2002). The transcriptional network initiated by Hcm1 is mediated in part via its transcriptional induction of *FKH* genes (Pramila *et al*. 2006); deletion of *HCM1* delayed G2/M progression and reduced cyclical *FKH* expression (Figs. 6A and 6B), as described previously (Pramila *et al.* 2006). Dividing *hcm1Δ* mutants displayed reduced *FKH* protein and mRNA levels (Figs. 6C and 6D), and as a consequence had impaired expression of *VMA* genes (Fig. 6E) and alkalized vacuoles (Fig. 6F) in young cycling cells. Hcm1 is therefore required for acidification of vacuoles, likely due to its induction of *FKH* genes.

**Figure 6.**
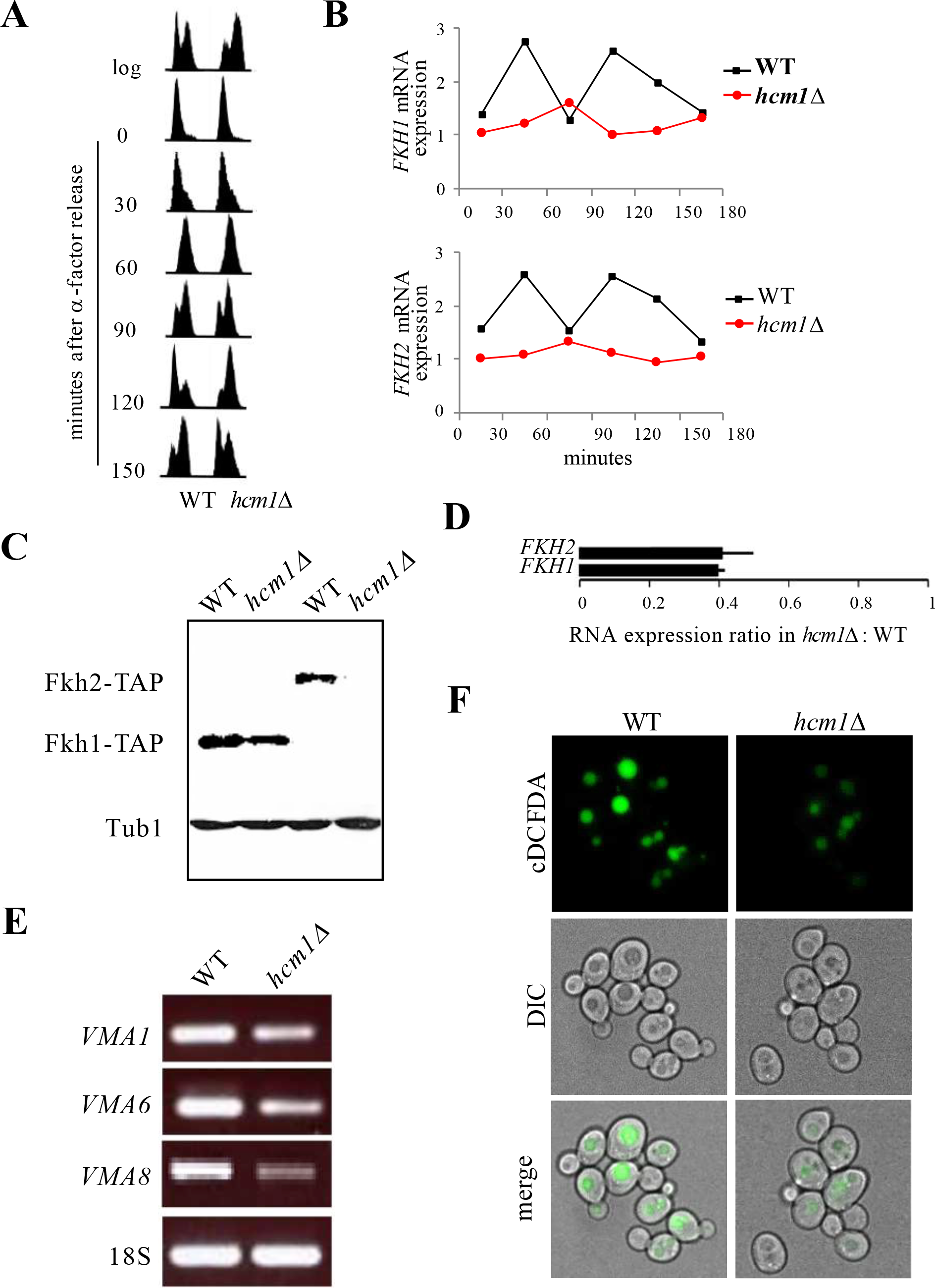
Hcm1 regulates *VMA* gene expression via upregulation of Fkh transcription. **(A)** Impaired progression through M phase in *hcm1Δ* mutants. Cells were arrested in G1 by α-factor treatment, released into fresh media, and analyzed by flow cytometry at the times indicated. **(B)** Cell cycle-dependent expression of *FKH1* and *FKH2* in yeast is lost in *hcm1*Δ cells. RNA was prepared from cells in (A) and analyzed by qPCR. Values were plotted to show changes over time. **(C)** TAP immunoblot of Fkh-TAP fusion proteins in whole cell lysates from *hcm1*Δ cells. Tubulin (Tub1) served as a loading control. **(D)** qPCR analyses of *FKH* mRNA expression in log *hcm1Δ* mutants relative to WT cells (mean ± SD). RNA extracted from cells used in (C). **(E)** Reduced expression of *VMA* genes in *hcm1Δ* mutants. RT-PCR was performed to assess *VMA* gene expression in *hcm1*Δ cells. **(F)** Live cell fluorescence images of dividing WT and *hcm1*Δ yeast cells stained with cDCFDA.

Hcm1 shuttles between the nucleus and the cytosol in a cell cycle dependent manner (Rodriguez-Colman *et al.* 2013). To determine whether Hcm1 nuclear-cytosolic shuttling is impaired in aging yeast cells, we assessed Hcm1-GFP cellular localization in dividing cells; while distributed throughout the cell during G2/M, Hcm1-GFP was exclusively nuclear during G1 (Fig. 7A). The cell cycle-dependent changes in Hcm1-GFP distribution were abolished in replicatively old unbudded G1 cells, compared to unbudded young cells, as Hcm1-GFP was no longer restricted to the nucleus (Fig. 7B). This was not a cell cycle-dependent affect, as budded and unbudded older cells all lacked nuclear Hcm1-GFP localization (Supplemental Fig. 5A). Fig. 7C confirmed that Hcm1 is indeed nuclear in young cells, and becomes rapidly excluded from nuclei in early budding (as early as 3 buds) G1 cells (representative images shown in Fig. 7C). Quantitation of the fluorescence intensity shows that the percent of Hcm1-GFP fluorescence localized to the nucleus falls as soon as mothers produce 3-4 buds (Fig. 7D). In contrast to Hcm1, Fkh1 and Fkh2 remain nuclear in young cycling cells (Supplemental Fig. 5B), and their localization was not influenced by the age of the cell (Fig. 7B; Supplemental Fig. 5A). These observations suggest that the cytosolic distribution of Hcm1 in old cells is not a general consequence of loss of nuclear integrity, but rather, the product of a specific impairment in shuttling Hcm1 into the nucleus. Impaired nuclear import of Hcm1 in early aging cells therefore provides a possible mechanism leading to reduced expression of the *FKH* genes, and the resultant loss of vacuolar acidity. Consistent with a downregulation in *FHK* and *VMA* mRNA, as well as loss of vacuolar acidity, *hcm1Δ* mutants were short lived compared to WT cells (Fig. 7E). Failure to induce *VMA* gene expression is likely a major contributor to reduced RLS in *hcm1*Δ cells, since constitutive expression of *VMA1* from a *GAL1* promoter partially restored RLS in these cells (Fig. 7E). Taken together, these results indicate that maintenance of vacuolar acidity, via upregulation of *VMA* gene expression, is important for enhanced RLS and is accomplished by Hcm1-dependent upregulation of *FKH1/FHK2* transcription (Fig. 8).

**Figure 7.**
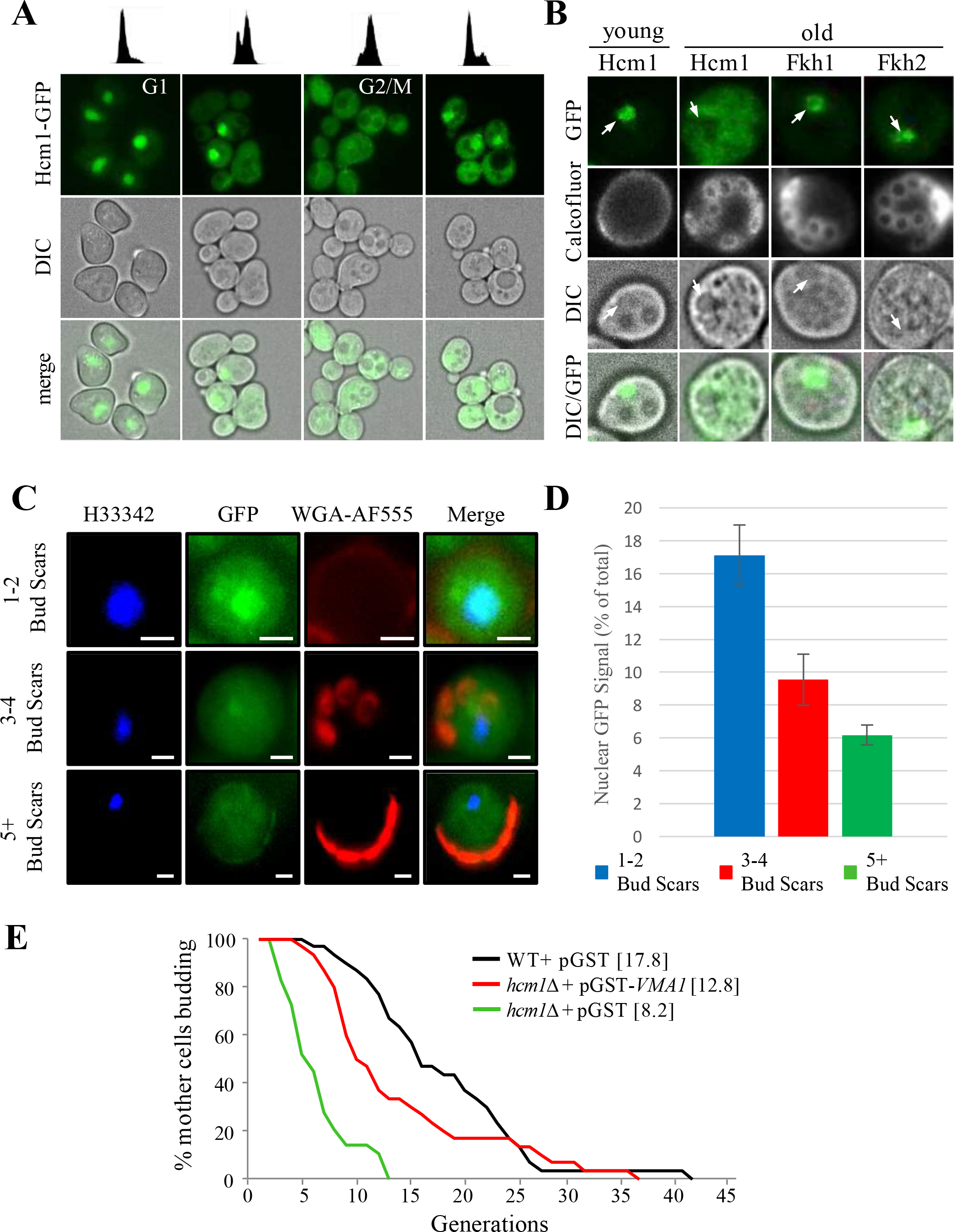
Nuclear exclusion of Hcm1 in replicatively old cells. **(A)** Nucleocytoplasmic distribution of Hcm1-GFP throughout the cell cycle. Cells were arrested in G1 by α-factor treatment, released into fresh media, and analyzed by flow cytometry at the cell cycle stage indicated (upper panel) and imaged (lower panels). Hcm1-GFP distribution in an asynchronous yeast culture is shown in (Supplemental Fig. 5B). **(B)** Live single cell fluorescence images of age-dependent Hcm1-GFP distribution. Arrows denote nuclei. Images of Fkh-GFP in age-matched cells are included for comparison. Calcofluor-white stain was added prior to imaging. See Supplemental Fig. 5A for additional images of older Fkh1-GFP, Fkh2-GFP and Hcm1-GFP expressing cells. **(C)** Live cell imaging of aging Hcm1-GFP cells. Nuclei were imaged using Hoechst (H33342; blue), bud scars were imaged using wheat germ agglutinin-alexafluor 555 conjugate (WGA-AF555; red) and Hcm1-GFP is shown in green (GFP). Cells were binned into cells with 1-2 bud scars, 3-4 bud scars and 5+ bud scars. All imaged cells were unbudded. 20 cells were image for the 1-2 BS group, 12 for the 3-4 BS group and 20 for the 5+ BS group. Representative images are shown. **(D)** Quantitation of GFP fluorescence intensity. The GFP signal in the cell is the sum of the pixel intensities across the whole cell while the nuclear signal is the sum of the GFP signal in each pixel inside the nucleus. The p-value for the 1-2 BS versus 3-4 BS groups was 0.002, the p-value for the 1-2 BS versus 5+ BS groups was 0.7 − 10Λ-5 (0.000007), and the p-value for the 3-4 BS versus 5+ BS groups was 0.04. **(E)** RLS analysis of WT cells and *hcm1Δ* mutants harboring the indicated plasmids. Cells were maintained on Ura^-^ synthetic media supplemented with 0.1% galactose to induce expression. Expression of *VMA1* partially restores the reduced RLS in *hcm1Δ* cells (p<0.001). Mean replicative lifespan shown in brackets (n=40).

**Figure 8.**
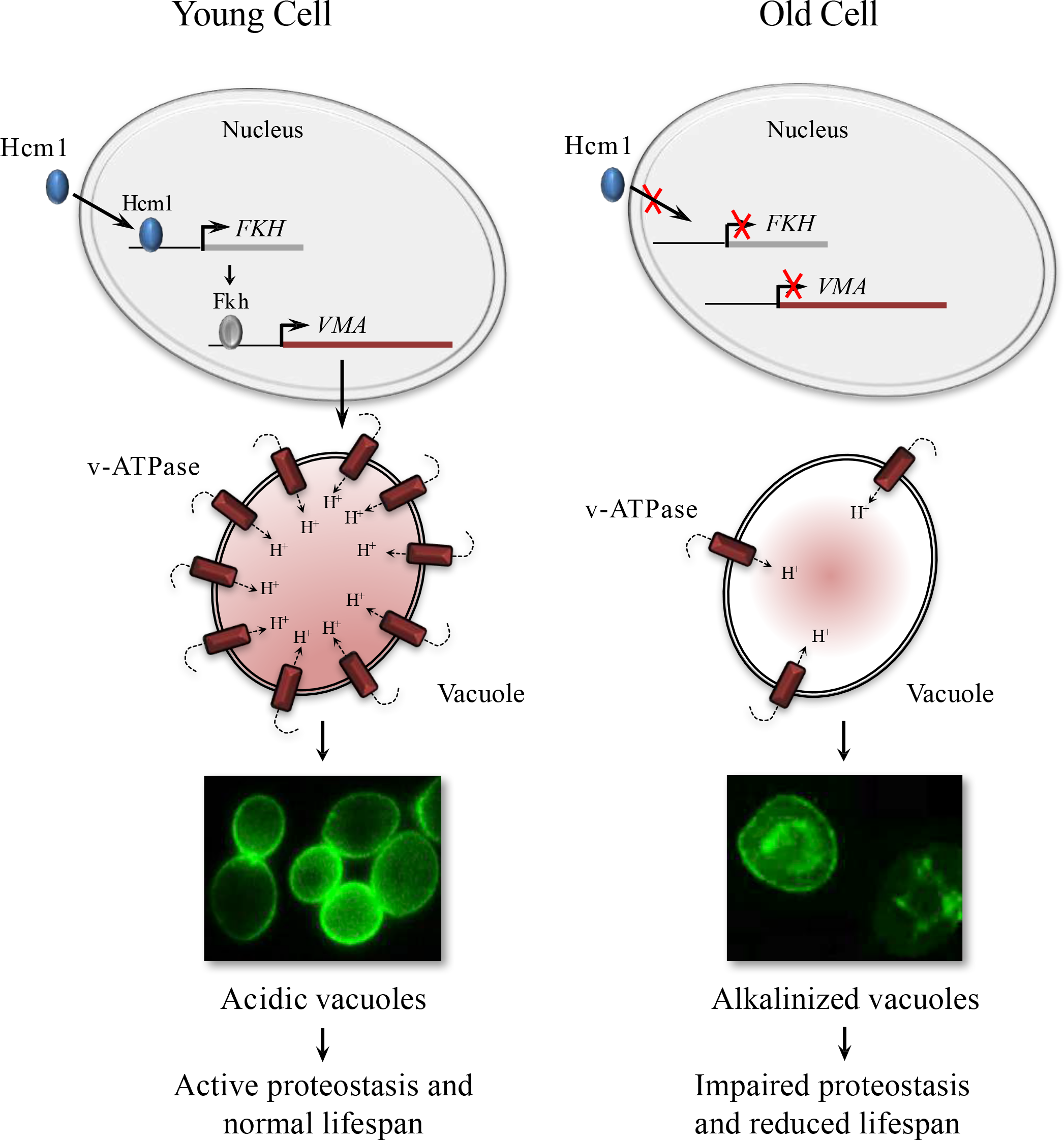
Schematic representation of control of vacuolar acidity in aging yeast cells. A working model of an interconnected transcriptional network underlying vacuolar acidity in yeast. Nuclear localization of Hcm1 upregulates the expression of *FKH* genes in S/G2, which, in turn, induces the expression of *VMA* vacuolar proton pump genes in cells undergoing early divisions. Hcm1 is no longer restricted to the nucleus in replicatively old cells. This results in reduced expression of *VMA* genes and the ensuing loss of vacuolar acidity.

### Discussion

Our work presented here describes a mechanism that regulates vacuolar acidity, proteostasis and replicative lifespan in yeast. Vacuolar acidity has now been shown by several reports, including the current report, to trigger replicative senescence when lost (Hughes and Gottschling 2012; Henderson *et al.* 2014; Baxi *et al.* 2017). Indeed, we show that transcription of vacuolar ATPase (v-ATPase) subunits, which constitute the assembly of a highly conserved enzyme responsible for acidifying vacuoles by pumping protons into the vacuolar lumen, is reduced as cells age (Fig. 2A). Vacuoles in dividing cells that lack v-ATPase subunits are alkalized, and these cells experience reduced replicative lifespan (RLS; Figs. 2B-2D). A compelling link between vacuolar acidity and longevity was identified when we demonstrated that v-ATPase subunit transcription is controlled by multiple members of the Forkhead Box (Fox) family of evolutionarily conserved stress responsive transcription factors (Fkh1, Fkh2 and Hcm1; Figs. 5 and 6). We observed that in mutants lacking both *FKH1* and *FKH2* vacuoles were alkalized, Fkh1 and Fkh2 both bound to select v-ATPase subunit promoters in WT cells, and subsequently, v-ATPase mRNA was reduced in *fkh1*Δ *fkh2*Δ cells. Interestingly, we also observed that *FKH1* and *FKH2* mRNA levels were reduced in older cells (Fig. 5D), providing a mechanism for why v-ATPase mRNA levels drop and vacuoles lose their acidity as cells age.

We had previously observed that cells lacking both *FKH1* and *FKH2* had reduced replicative and chronological lifespan (Postnikoff *et al.* 2012). At the time, the Fkh proteins had not been linked with vacuolar function, but our report was the first to show that Fkh-dependent longevity was conserved in yeast, as it was also apparent in higher eukaryotic models (Murphy and Hu 2013; Martins *et al.* 2016). At the time, we connected the Fkh proteins with the yeast Anaphase Promoting Complex (APC) in a stress responsive capacity that was required for cell health. Stress response also involves autophagy, which utilizes vacuoles to rid the cell of unfolded proteins and protein aggregates (Cuervo *et al.* 2005; Tang *et al.* 2008), and we showed that the induction of enhanced yeast RLS as a result of ER stress is due to elevated autophagy (Ghavidel *et al.* 2015). Although the Fkh proteins and the APC work together to alleviate stress in yeast cells, we do not currently know whether the APC impacts autophagy and vacuolar acidity. It can thus be envisioned that maintenance of vacuolar acidity is a critical component of an interconnected stress response network.

The third member of the Fox family we identified in this study is Hcm1. Hcm1, Fkh1 and Fkh2 are part of temporally regulated cell cycle progression pathway; Hcm1 is transcribed during G1/S and then transcribes the Fkhs during G1 (Horak *et al*. 2002; Pramila *et al.* 2006). In our studies to determine why the Fkh proteins are reduced in old cells, we observed that Hcm1 cellular distribution is altered in old versus young cells (Figs. 7B-D). While the Fkhs remained nuclear throughout the cell cycle, which did not change in young or old cells (Figs. 7B and 7D; Supplemental Fig. 5), Hcm1 is exclusively nuclear in young unbudded G1 cells, but distributed throughout the cell during G2/M (Fig. 7A). At early stages of aging (3 bud stage), on the other hand, Hcm1 is excluded from the nucleus in unbudded G1 cells (Figs. 7C, D). Indeed, Hcm1 is excluded from the nucleus in all phases of the cell cycle in older cells (Supplemental Fig. 5). As predicted from this observation, vacuoles are alkalized, v-ATPase subunit mRNA levels are reduced, and RLS is decreased in *hcm1*Δ cells (Figs. 6 and 7). This provides a mechanism whereby Hcm1 nuclear shuttling is stalled as cells age, resulting in reduced Fkh function, and replicative senescence. Like the Fkhs, in addition to cell cycle control, Hcm1 is also involved in the stress response (Rodriguez-Colman *et al.* 2013; Maoz *et al.* 2015). In response to stress, the stress response kinase Snf1 phosphorylates Hcm1, driving it into the nucleus to initiate its transcriptional program (Rodriguez-Colman *et al.* 2013). This defines a further player in our proposed interconnected stress response network.

The reason Hcm1 nuclear import is blocked as cells age remains unresolved. Hcm1 was recently shown to be required for RLS in a caloric restriction independent manner (Maoz *et al.* 2015). In retrospect, this would be predicted based on observations presented here. We recently found that Hcm1 is required for the induction of Snf1 kinase activation in response to stress (Jiao *et al.* 2016). This mechanism, like the one presented here, required the transcriptional activation of the *FKH* genes in response to stress, which then went on to transcribe *SNF1*. In that report, we also found that deletion of the *UBC1* ubiquitin conjugating enzyme impaired Snf1 function, and that this was due to impaired nuclear shuttling of Hcm1. Subsequently, we found that cells lacking *UBC1* have reduced chronological lifespan (Harkness, unpublished data), consistent with its role in activating the Fox stress response and lifespan extending pathway (Jiao *et al.* 2016). The impact Ubc1 has on Hcm1 is yet to be determined, as Hcm1 protein levels are reduced in *ubc1*Δ cells, but its stability is not altered, indicating that Ubc1-dependent degradation of Hcm1 may not occur (Jiao *et al.* 2016). *HCM1* itself is expressed in a cyclical manner in G1/S by the Swi4/Swi6 complex (Pramila *et al.* 2006). It remains possible, but untested, that Ubc1 alters Swi4/Swi6 function. Alternatively, phosphorylation of Hcm1 by Snf1 (Rodriguez-Colman *et al.* 2013) may perhaps make Hcm1 a target for Ubc1 mediated (mono)ubiquitination (Rodrigo-Brenni and Morgan 2007). Hcm1 modified in this manner may then be capable of transiting the nuclear membrane, as previously described for other shuttling proteins (Kragt *et al.* 2005). Future work will be dedicated to assessing these possibilities. Thus, the stress response network described here likely requires input from the ubiquitin-signalling cascade, which regulates Hcm1 nuclear shuttling in an as yet uncharacterized manner.

Our data strongly supports the idea that reduced RLS in response to vacuolar alkalization occurs at least in some part through impaired proteostasis. Hydrolases within vacuoles require an acidic pH to function, thus loss of vacuolar acidity should cause loss of hydrolase function and protein aggregation (Hecht *et al.* 2014). Increased protein aggregation is linked with aging and many aging related neurological diseases, while enhanced proteostasis increases lifespan in model organisms (Noormohammadi *et al.* 2016; Cuanalo-Contreras *et al.* 2017; Currais *et al.* 2017). Thus, loss of proteostasis is predicted to be a primary source of aging in older yeast cells with alkalized vacuoles. However, the specific events that connect vacuolar alkalization to replicative senescence remain obscure, although there is evidence to suggest mitochondrial dysfunction may be linked to the phenomena (Ohya *et al.* 1991; Merz and Westermann 2009; Hughes and Gottschling 2012). The potential loss of mitochondria function upon vacuolar impairment is consistent with recent thought on the role of intracellular organelle communication in cell function (Lang *et al.* 2015; Honscher and Ungermann 2014; Dakik and Titorenko 2016). To test whether protein aggregation was involved in replicative senescence, we used the human α-synuclein protein tagged with GFP (α-syn-GFP) as a surrogate marker for the accumulation of protein inclusions. α-syn is a protein found inappropriately folded and aggregated in human neurological diseases (Goedert *et al.* 2017). We indeed observed increased accumulation of α-syn-GFP in older yeast cells, and in cells lacking the genes encoding the v-ATPase subunit Vma8 or the vacuolar protease Pep4 (Figs. 3A-3C). Thus, as expected, α-syn-GFP was observed to aggregate and accumulate in cells with impaired vacuolar function. Finally, reduced RLS in cells expressing α-syn-GFP (Fig. 4D) strongly supports the idea that vacuolar proteostasis plays a pivotal role in maintaining yeast replicative lifespan.

Taken together, the results presented in this report describe an interconnected stress response pathway that maintains vacuolar pH in young cells, but vacuolar pH erodes as yeast cells age (Fig. 8). In a parallel study (Baxi *et al.* 2017), we describe the loss of lysosomal acidity in the intestine of post-reproductive *C. elegans* and implicated gonad to soma signalling in its regulation. As in yeast, *C. elegans* recruits a Forkhead protein (DAF-16) to upregulate v-ATPase gene transcription and acidify lysosomes, preventing protein aggregation and premature senescence during early adulthood, the lifecycle stage of maximal reproduction. As animals leave reproductive life, a time coincident with the start of protein aggregation in the soma, DAF-16 is excluded from the nucleus. These studies reveal a remarkable conservation in a cellular mechanism coupling protein catabolism and proliferative capacity and suggest its early co-optation in metazoans to mediate germline to soma communication (Baxi *et al.* 2017). Why the Fox network fails in older yeast and worms remains unexplained, but if mechanisms can be discovered that extend Fox function over time, it should be possible to enhance proteostasis, and promote longevity.

## Acknowledgements

This work was supported by grants to TAH by CIHR, NSERC and CFI, to CEC by NSERC and CFI, and to CHE by NSERC.

## Supplemental Figure Legends

**Figure S1.**
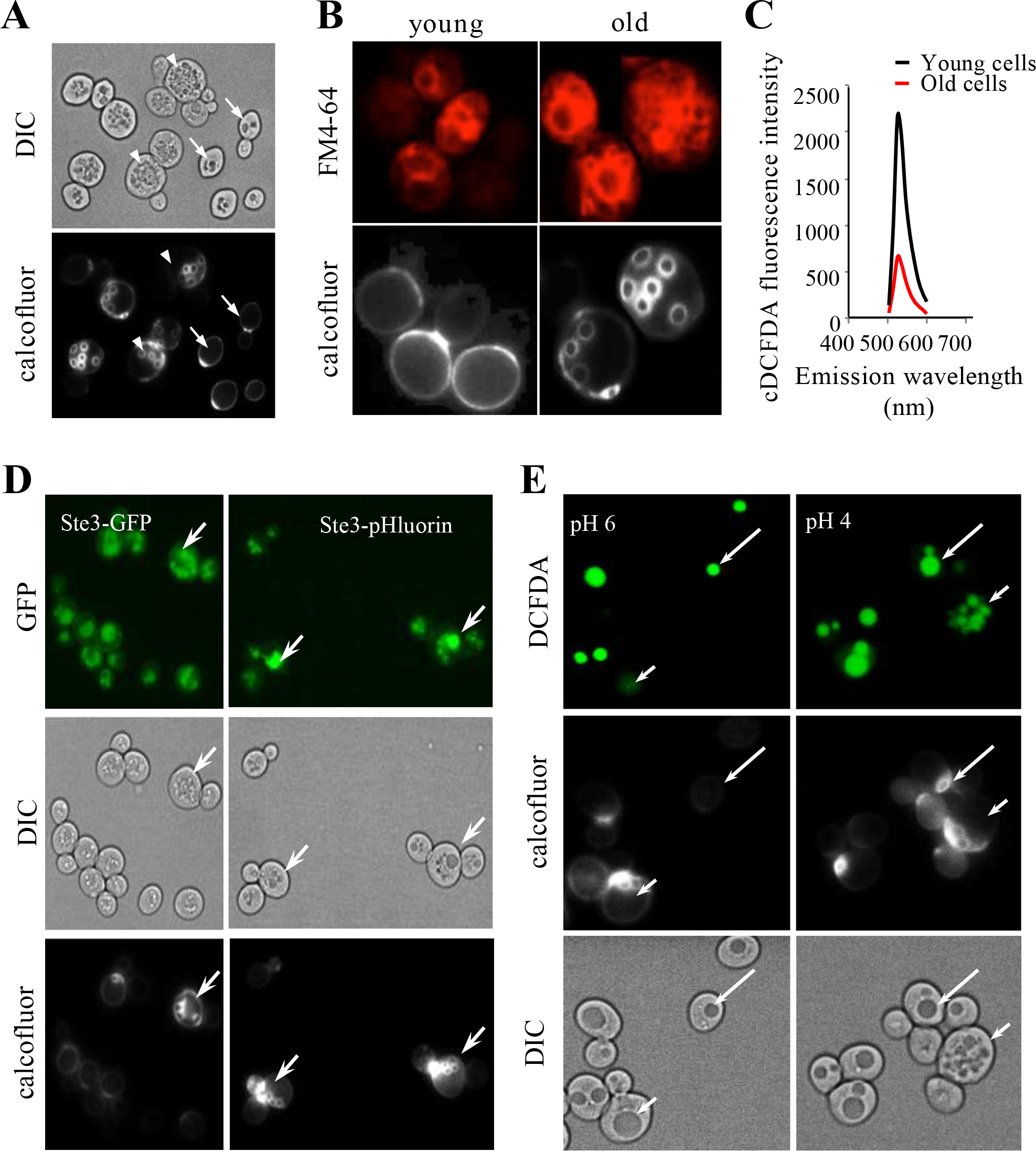
**(A)** Live cell imaging of dividing WT yeast. Bud scars were stained with calcofluor-white prior to imaging. Arrows and arrowheads denote representative young and old cells, respectively. (**B**) Live cell imaging of dividing yeast co-stained with the vacuolar membrane marker FM4-64 and calcofluor-white. (**C**) cDCFDA fluorescence emission spectra of young and old yeast. Spectra from young cells with less than 2 bud scars and old cells with 12-20 bud scars are plotted (n=20). **(D)** Live cell imaging of dividing Matα yeast cells expressing either Ste3-GFP, or the GFP variant pHluorin fused to the pheromone receptor Ste3. Ste3-GFP and Ste3-pHluorin are both targeted to yeast vacuoles. pHluorin fluorescence becomes visible as the pH of the local environment increases. Arrows indicate older cells. **(E)** Following overnight growth in standard YPD media (pH 6.0), yeast were resuspended in acidic (pH 4.0) YPD media and grown for an additional 6 hours to acidify vacuoles. The lipid soluble pH sensitive dye cDCFDA was then added to cultures, which will fluoresce when exposed to acidic environments. Long arrows denote young cells, while short cells identify older cells.

**Figure S2.**
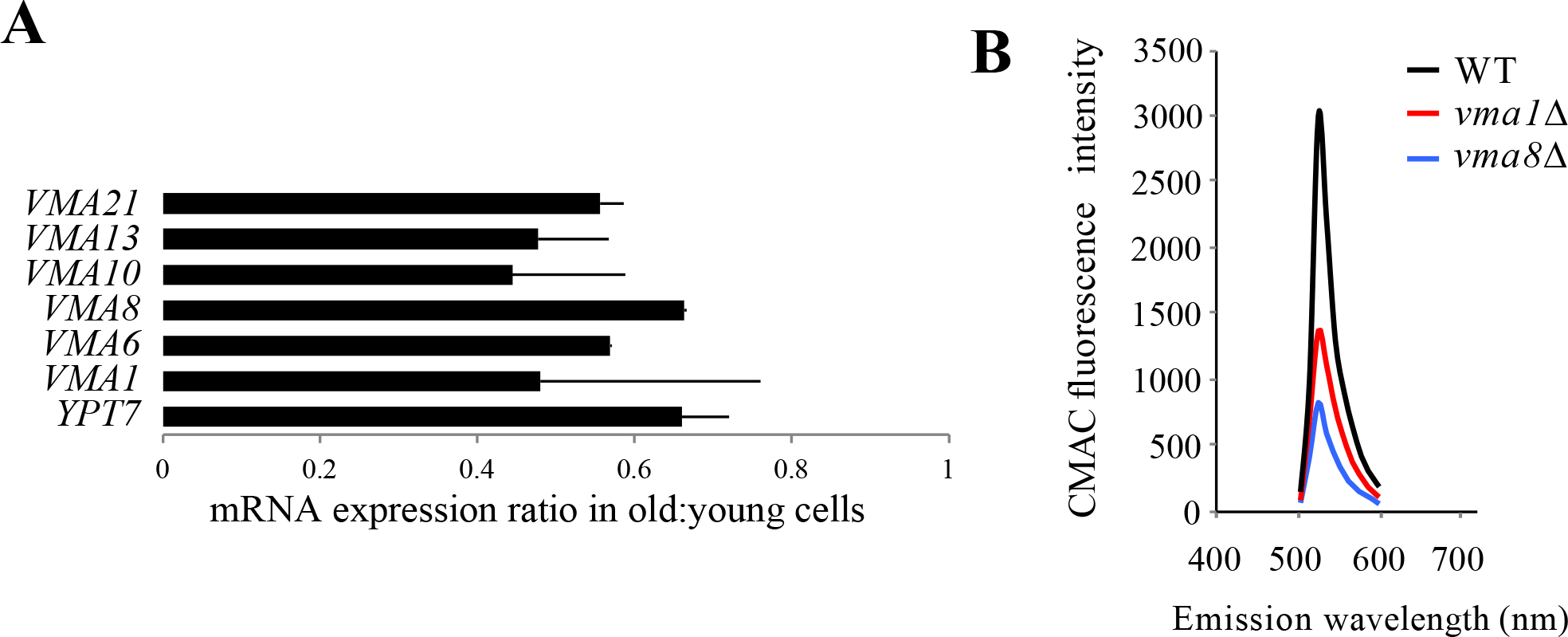
**(A)** Expression ratios of RT-PCR RNA expression analyses of vacuolar proton pump subunits (*VMA*) and a vacuolar fusion factor (*YPT7*) in replicatively old cells enriched using MEP. Expression ratios were normalized to 18S rRNA and plotted. Mean ± SD of three individual analyses from two RNA preparations are shown. **(B)** CMAC-Arg fluorescence emission spectra of cells in (Fig. 2B) (n=25).

**Figure S3.**
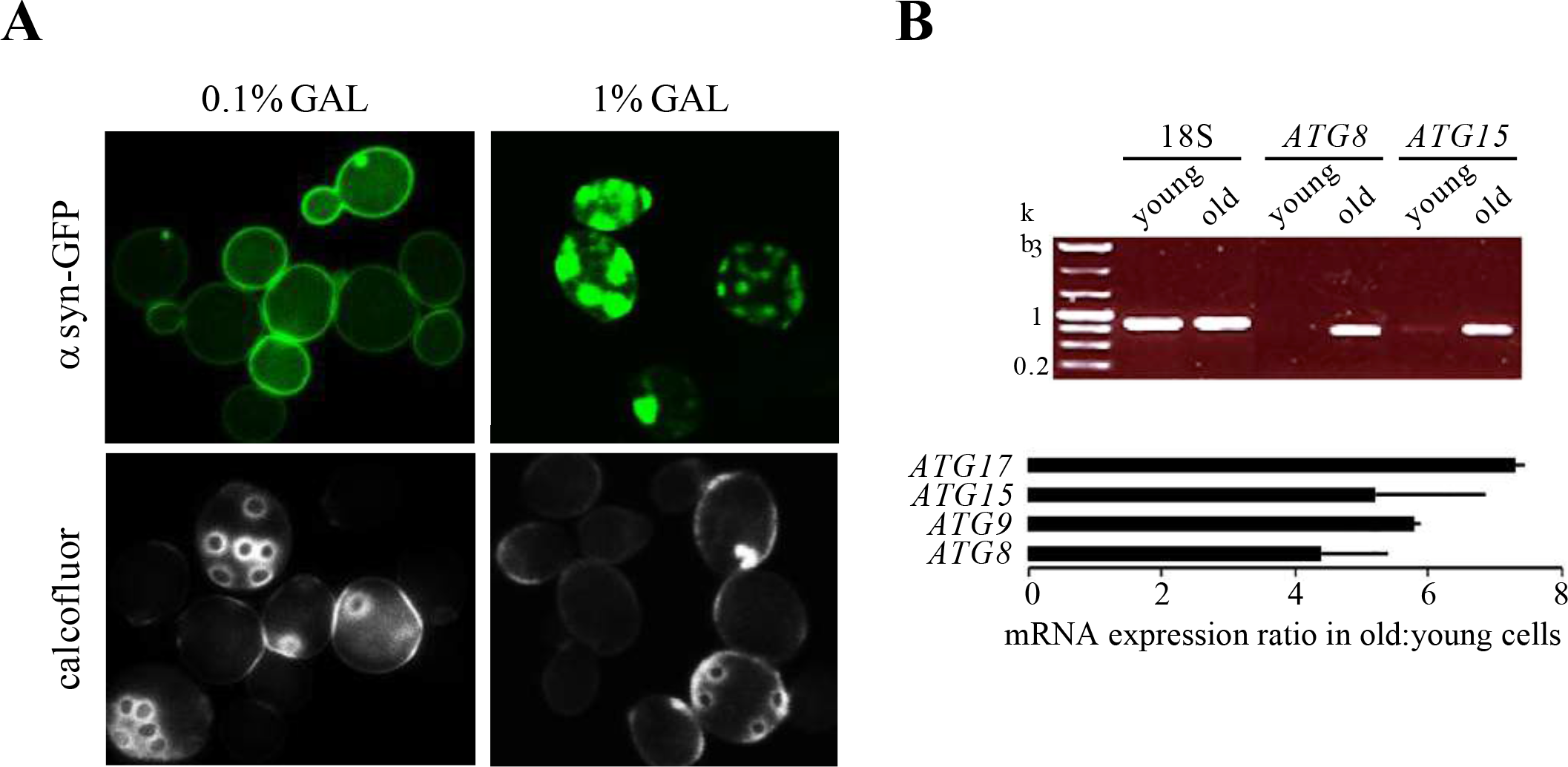
**(A)** Live cell microscopy of WT yeast expressing a genomically integrated gal-inducible α-syn-GFP construct. Cells were counterstained with calcofluor before imaging. A further increase in α-syn-GFP expression in 1% galactose resulted in increasingly larger α-syn-GFP aggregates. **(B)** RNA expression analysis of autophagosome (ATG) subunits in early log (young) cells and a population enriched for replicatively old cells using MEP. Upper panel shows agarose gel electrophoresis of ethidium bromide stained RT-PCR products. RNA expression ratios in old:young cells determined by qPCR were normalized to 18S rRNA and plotted in the bottom panel. Mean ± SD of three individual analyses from two RNA preparations are shown.

**Figure S4.**
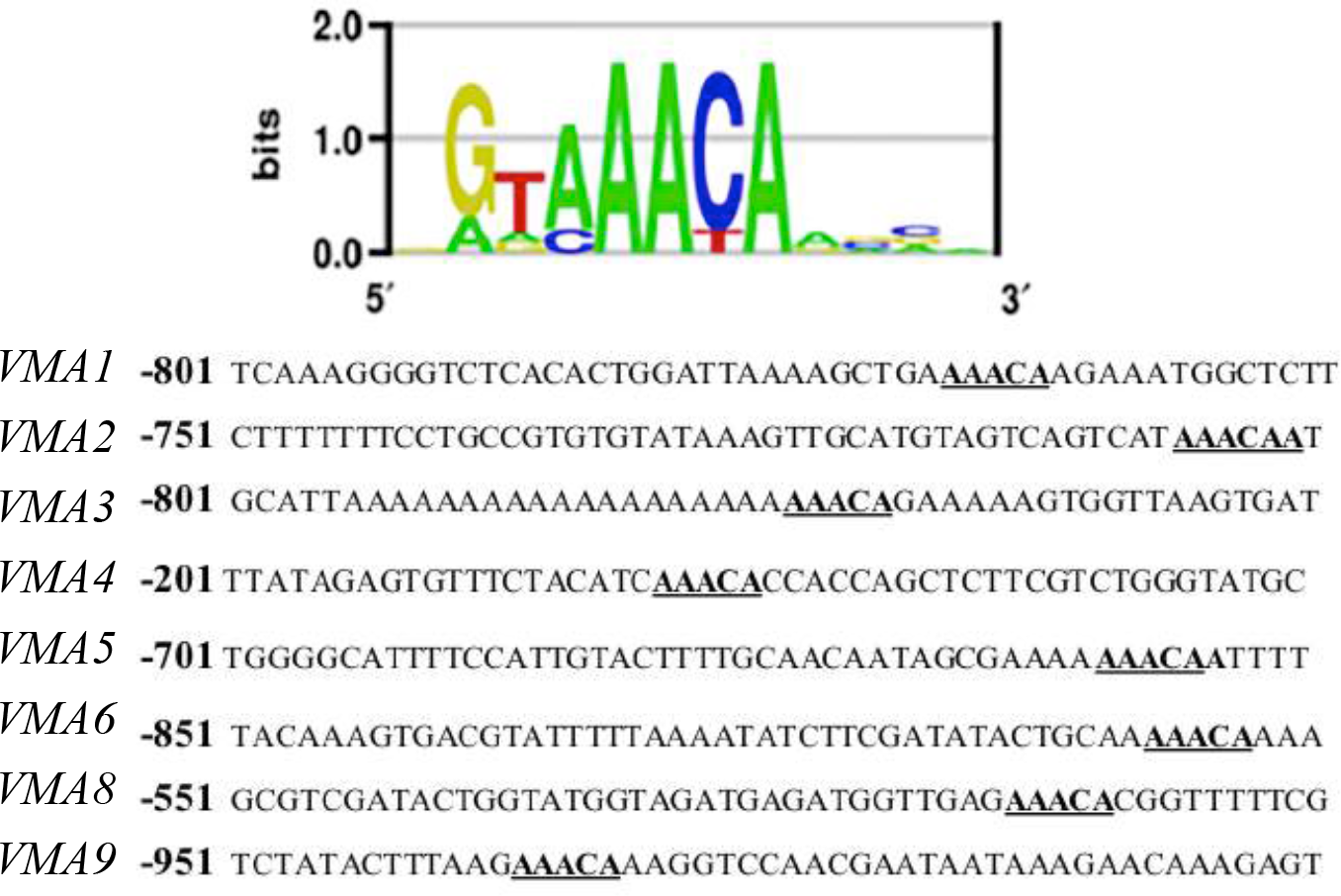
Consensus binding motifs for Forkhead transcription factors in the 5’ upstream regions of *VMA* genes (http://yetfasco.ccbr.utoronto.ca). Core binding domains are bolded and underlined.

**Figure S5.**
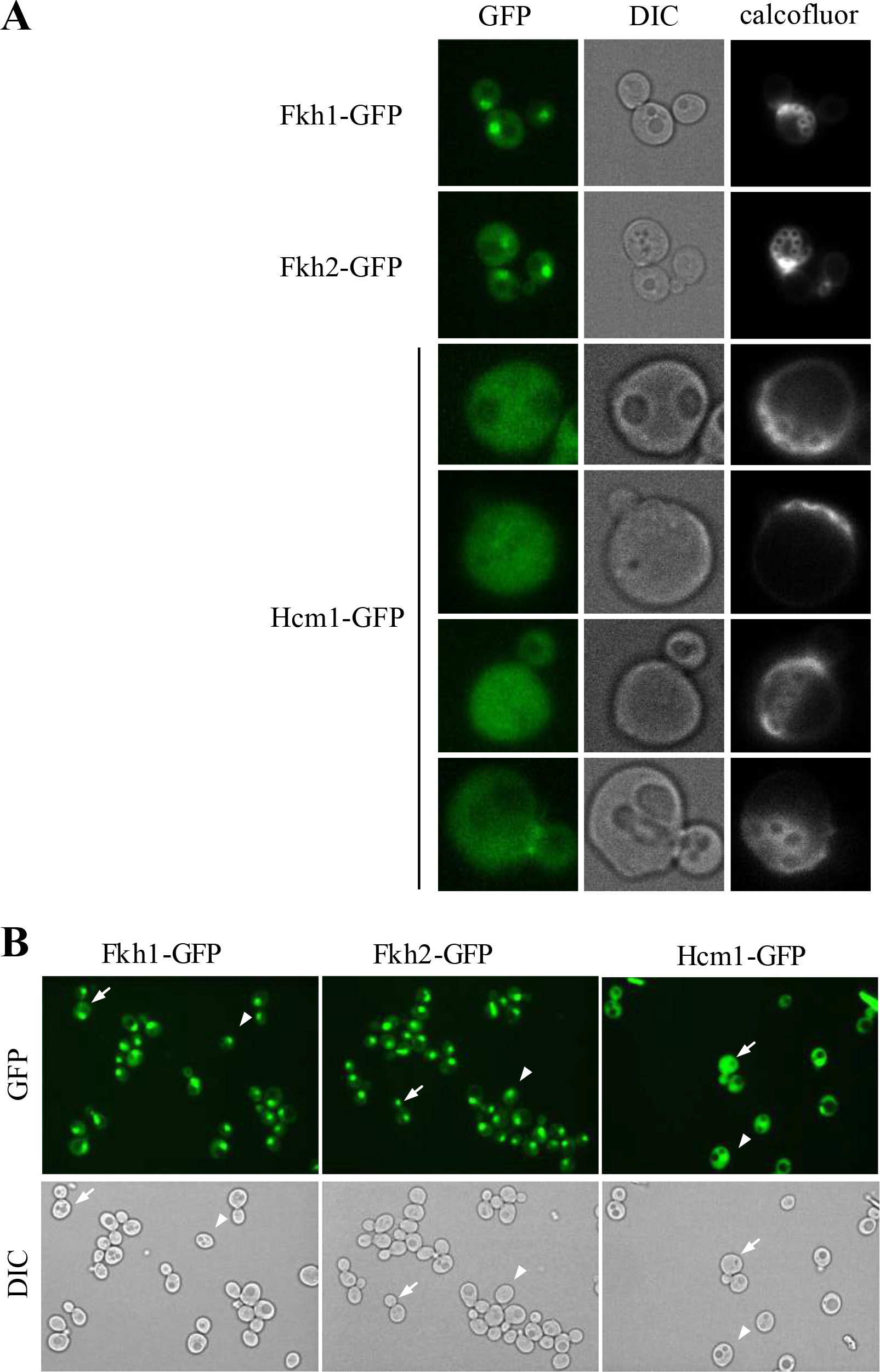
Additional images of aging cells expressing endogenous GFP C-terminal fusions to the *FKH1*, *FKH2* and *HCM1* genes. **(A)** Calcofluor was added prior to imaging. Fkh1 and Fkh2 remain nuclear throughout the cell cycle and in aging cells. Hcm1, on the other hand, is excluded from the nucleus in aging cells. **(B)** Live cell fluorescence microscopy of GFP fusions in dividing yeast cultures. Arrows and arrowheads denote cells in G2/M and G1, respectively. Hcm1 is nucleocytoplasmic in budded cells (G2/M), but restricted to the nucleus in G1 cells. As seen above, Fkh proteins maintain a nuclear localization throughout the cell cycle.

---

**Table S1.** Statistical analysis for RTLS of vma1Δ mutants (Fig. 2D).

|  | *vma1* | *vma8* | *ypt7* | WT |
| --- | --- | --- | --- | --- |
| T-test (compared with wt) | 0.0032 | 9.64E-05 | 0.10 | - |
| Mann-Whitney Test (compared to wt) | 0.0403 | 1.75E-04 | 0.10 | - |
| Significant difference from wt (p < 0.05) | Yes | Yes | No | - |
| Mean Lifespan (nmd*) | 13.25 | 10.63 | 17.83 | 20.50 |
| Maximal Lifespan (top 10%, nmd) | 17.45 | 14.30 | 24.50 | 40.05 |

\* nmd – number of mitotic divisions

**Table S2.** Cytoplasmic and nuclear GFP signal in cells with 1-2 bud scars.

| N | Cytoplasmic Area (Pixels) | Total Cytoplasmic GFP Signal | Nuclear Area (Pixels) | Total Nuclear GFP Signal | Percent Nuclear GFP |
| --- | --- | --- | --- | --- | --- |
| 1 | 1150 | 114971 | 63 | 9260 | 8.05 |
| 2 | 1432 | 140398 | 246 | 31934 | 22.75 |
| 3 | 1478 | 103989 | 174 | 19359 | 18.62 |
| 4 | 1223 | 135378 | 114 | 21234 | 15.69 |
| 5 | 1261 | 149667 | 100 | 19009 | 12.70 |
| 6 | 2445 | 306720 | 383 | 67458 | 21.99 |
| 7 | 1980 | 159408 | 199 | 26071 | 16.35 |
| 8 | 1855 | 189781 | 337 | 53669 | 28.28 |
| 9 | 2316 | 196770 | 339 | 45405 | 23.08 |
| 10 | 1894 | 144688 | 113 | 13751 | 9.50 |
| 11 | 2516 | 282001 | 384 | 59480 | 21.09 |
| 12 | 1018 | 181786 | 340 | 75220 | 41.38 |
| 13 | 2346 | 316780 | 299 | 66492 | 20.99 |
| 14 | 2305 | 248896 | 160 | 29970 | 12.04 |
| 15 | 2415 | 323967 | 299 | 66492 | 20.52 |
| 16 | 2588 | 293826 | 246 | 42684 | 14.53 |
| 17 | 2219 | 188074 | 78 | 12824 | 6.82 |
| 18 | 2358 | 238505 | 123 | 19057 | 7.99 |
| 19 | 1990 | 284520 | 163 | 37912 | 13.32 |
| 20 | 1450 | 133631 | 60 | 8891 | 6.65 |

**Table S3.** Cytoplasmic and nuclear GFP signal in cells with 3-4 bud scars.

| N | Cytoplasmic Area (Pixels) | Total Cytoplasmic GFP Signal | Nuclear Area (Pixels) | Total Nuclear GFP Signal | Percent Nuclear GFP |
| --- | --- | --- | --- | --- | --- |
| 1 | 2493 | 244374 | 267 | 32318 | 13.22 |
| 2 | 3380 | 379655 | 454 | 77361 | 20.38 |
| 3 | 2965 | 414534 | 381 | 69748 | 16.83 |
| 4 | 2829 | 248457 | 287 | 36334 | 14.62 |
| 5 | 2454 | 211792 | 172 | 17295 | 8.17 |
| 6 | 2512 | 192040 | 126 | 10861 | 5.66 |
| 7 | 2184 | 180562 | 72 | 7539 | 4.18 |
| 8 | 1669 | 124886 | 43 | 4193 | 3.36 |
| 9 | 2280 | 304644 | 146 | 26380 | 8.66 |
| 10 | 3473 | 305624 | 76 | 8262 | 2.70 |
| 11 | 3016 | 194065 | 149 | 13277 | 6.84 |
| 12 | 3502 | 340647 | 203 | 33624 | 9.87 |

**Table S4.** Cytoplasmic and nuclear GFP signal in cells with 5+ bud scars.

| N | Cytoplasmic Area (Pixels) | Total Cytoplasmic GFP Signal | Nuclear Area (Pixels) | Total Nuclear GFP Signal | Percent Nuclear GFP |
| --- | --- | --- | --- | --- | --- |
| 1 | 2084 | 274575 | 216 | 17014 | 6.20 |
| 2 | 2642 | 284234 | 97 | 10566 | 3.72 |
| 3 | 3618 | 328938 | 117 | 11870 | 3.61 |
| 4 | 3340 | 264942 | 186 | 20844 | 7.87 |
| 5 | 3011 | 318145 | 253 | 36211 | 11.38 |
| 6 | 2593 | 192110 | 126 | 10770 | 5.61 |
| 7 | 3755 | 289785 | 142 | 16313 | 5.63 |
| 8 | 2685 | 182808 | 128 | 10523 | 5.76 |
| 9 | 2704 | 289247 | 248 | 31692 | 10.96 |
| 10 | 3482 | 234077 | 100 | 9420 | 4.02 |
| 11 | 3827 | 454908 | 229 | 27401 | 6.02 |
| 12 | 3512 | 245496 | 159 | 13179 | 5.37 |
| 13 | 4406 | 362927 | 225 | 25825 | 7.12 |
| 14 | 3405 | 352309 | 100 | 12918 | 3.67 |
| 15 | 2998 | 168308 | 54 | 4395 | 2.61 |
| 16 | 2184 | 180562 | 72 | 7539 | 4.18 |
| 17 | 3563 | 279232 | 202 | 23035 | 8.25 |
| 18 | 2602 | 179379 | 128 | 10523 | 5.87 |
| 19 | 2812 | 163765 | 73 | 5511 | 3.37 |
| 20 | 2063 | 141996 | 193 | 17636 | 12.42 |

**Table S5.** Summary of Hcm1-GFP imaging statistics.

| Group | Average % Nuclear GFP | SEM | P-Value Versus 0 - 2 BS Group | P-Value Versus 3 - 4 BS Group |
| --- | --- | --- | --- | --- |
| 0 - 2 BS | 17.12 | 1.84 | N/A | 0.00246 |
| 3 - 4 BS | 9.54 | 1.56 | 0.00246 | N/A |
| 5+ BS | 6.18 | 0.61 | 0.00001 | 0.04 |

